# Single-cell and bulk transcriptional profiling of mouse ovaries reveals novel genes and pathways associated with DNA damage response in oocytes

**DOI:** 10.1101/2024.02.02.578648

**Authors:** Monique Mills, Chihiro Emori, Parveen Kumar, Zachary Boucher, Joshy George, Ewelina Bolcun-Filas

## Abstract

Immature oocytes enclosed in primordial follicles stored in female ovaries are under constant threat of DNA damage induced by endogenous and exogenous factors. Checkpoint kinase 2 (CHEK2) is a key mediator of the DNA damage response in all cells. Genetic studies have shown that CHEK2 and its downstream targets, p53 and TAp63, regulate primordial follicle elimination in response to DNA damage, however the mechanism leading to their demise is still poorly characterized. Single-cell and bulk RNA sequencing were used to determine the DNA damage response in wildtype and *Chek2*-deficient ovaries. A low but oocyte-lethal dose of ionizing radiation induces a DNA damage response in ovarian cells that is solely dependent on CHEK2. DNA damage activates multiple ovarian response pathways related to apoptosis, p53, interferon signaling, inflammation, cell adhesion, and intercellular communication. These pathways are differentially employed by different ovarian cell types, with oocytes disproportionately affected by radiation. Novel genes and pathways are induced by radiation specifically in oocytes, shedding light on their sensitivity to DNA damage, and implicating a coordinated response between oocytes and pre-granulosa cells within the follicle. These findings provide a foundation for future studies on the specific mechanisms regulating oocyte survival in the context of aging, as well as therapeutic and environmental genotoxic exposures.

## Introduction

DNA damage response (DDR) plays an essential role in cell and organ function. DNA damage triggers the DDR, a complex network of cellular pathways that detect, signal, repair the damage, and trigger apoptosis if damage is irreparable. However, the response mechanisms and cellular outcomes of DNA damage may differ depending on the cell type including cell death, senescence, or terminal differentiation (1–4). DNA damage encountered by cells via endogenous and exogenous mechanisms is also implicated in cellular and organ aging (5–7). The ovaries are organs that naturally age early due to a lack of regenerative abilities, resulting in cessation of estrogen production and menopause at the average age of 50-51 (8). Estrogen is produced by growing follicles recruited from the primordial follicle (PF) pool. Each female is born with a finite number of PFs, each containing an immature oocyte surrounded by somatic cells (primordial oocyte). As a woman ages, the PF reserve gradually diminishes, and the remaining PFs become more susceptible to DNA damage and other forms of cellular stress. Endogenous DNA damage may arise during normal cellular activities such as replication, transcription, or result from reactive oxygen species (ROS) produced during normal metabolic processes. Exogenous DNA damage can be caused by environmental exposures such as ionizing radiation (IR), alkylating agents, heavy metals, and various chemicals. For ovaries, it has been shown that IR- and chemotherapy-induced DNA damage causes accelerated depletion of the ovarian follicle reserve (9,10). Multiple studies have highlighted the critical role of checkpoint kinase 2 (CHEK2) in the establishment and maintenance of the ovarian follicle reserve (11–15). Moreover, *CHEK2* loss-of-function variants in humans correlate with females having a larger ovarian follicle reserve and later onset of menopause (16,17). These indicate that CHEK2-signaling in the ovary regulates follicle survival during women’s reproductive life not only in response to genotoxic insults, but also in response to endogenous sources of DNA damage. Improper repair of DNA damage can lead to mutations and genomic rearrangements. Therefore, cells have evolved a DDR and quality checkpoints that ensure DNA fidelity before cells can divide. DNA double strand breaks (DSBs) are the most lethal type of DNA damage. Cells respond to DSBs by activating a multi-layered signaling cascade involving many proteins and processes and CHEK2 is a key checkpoint kinase mediating DDR (18). CHEK2 phosphorylates and activates TRP53 (henceforth p53), which in turn plays a critical role in coordinating DDR outcomes and cell fate decisions, such as cell cycle arrest or apoptosis. Cell cycle arrest allows more time to repair DSBs, after which cells can re-enter the cell cycle. If DNA damage persists unrepaired, p53 triggers apoptosis. In contrast to somatic cells, which rely mainly on p53 activity, oocytes express a related protein TRP63 (p63) (19,20). TA isoform of p63 (henceforth TAp63) expressed exclusively in oocytes in the ovary has been shown to trigger oocyte apoptosis after activation by CHEK2 in response to DNA damage (11,12,21). Animal studies show that inactivation of CHEK2 or TAp63 but not p53, prevents apoptotic elimination of oocytes exposed to low doses of IR (0.2 - 0.5 Gy) (11,19,20). This indicates that for primordial oocytes—whose continuous meiotic arrest (months in mice and decades in humans) could allow for efficient repair—the primary response to DSBs appears to be an imminent apoptotic elimination. However, our knowledge of DDR in primordial oocytes is still minimal compared to somatic cells or fully grown oocytes and mostly based on the activities of the few aforementioned proteins (22–24).

To further our understanding of the mechanism regulating primordial oocyte survival or death in response to genotoxic insults, we conducted bulk and single-cell transcriptomic analysis of ovarian response to DNA damage in wildtype and CHEK2-deficient females. To model DNA damage, we employed IR, which is known to induce DSBs and apoptosis in primordial oocytes. We show that—in addition to apoptosis—DNA damage activates multiple response pathways in the ovary related to p53, interferon signaling, inflammation, cell adhesion and intercellular communication. Our results indicate that different cell types within the ovary employ these pathways differently and that oocytes mount the strongest cellular response. We identified novel genes involved in oocyte-specific DDR that may contribute to primordial oocyte sensitivity to DNA damage. Considering the importance of DNA damage in the process of aging, exposure to chemicals, and genotoxic therapies, our findings could pave the way for treatment strategies aimed at delaying ovarian aging and mitigating the toxic side effects of anti-cancer treatments.

## Results

### Radiation-induced DNA damage in ovaries activates CHEK2 dependent signaling

A single dose of relatively low radiation (≤0.5 Gy) eradicates primordial oocytes in mice within a few days. In contrast, growing oocytes (those inside growing follicles) persist longer (**Figure 1A**). The elimination of primordial oocytes is dependent on CHEK2 as abundant primordial oocytes are present in *Chek2-/-* ovaries after radiation (**Figure 1B**). IR induces DNA DSBs, which were readily detectable in ovaries after IR; all oocytes stained positive for DNA damage marker γH2AX (**Figure 1C**). IR-induced damage activated CHEK2 as evidenced by the positive staining for phosphorylated CHEK2 at threonine 68 (pCHEK2)(25) (**Figure 1D**). Similar to γH2AX, pCHEK2 was predominantly activated in oocytes (**Figure 1D**). This suggests that IR-induced damage triggers a different response in oocytes than in pre-granulosa cells or other ovarian somatic cells. In addition to direct DNA damage, IR also induces oxidative damage (26,27). The most common oxidative DNA lesion, 8-oxo-2′-deoxyguanosine (8-oxo-dG), was detected after IR in both oocytes and pre-granulosa cells of the primordial follicle (**Figure 1E**). These results confirm that IR induces direct DNA damage and oxidative damage which activates the DDR in the ovary. CHEK2 phosphorylates p53 in response to DSBs, which prevents p53 degradation by an MDM2-dependent mechanism (28). Because p53 seems dispensable for oocyte apoptosis, we next tested p53 activation by IR in ovaries. Phosphorylated p53 (S15) was detected in oocytes, pre-granulosa cells, and other cell types in irradiated ovaries (**Figure 1F**). TAp63 is a direct target of CHEK2 phosphorylation and the key proapoptotic factor in oocytes (11,12,20). Therefore, activation of p53 in oocytes after IR suggests that p53 may still play a role in the DDR and potentially regulates other cellular responses in oocytes. To summarize, IR exposure induces direct and indirect DNA damage in oocytes—and to a lesser extent in somatic cells—leading to activation of CHEK2-dependent signaling through TAp63 and p53. In the absence of CHEK2, TAp63 and p53 fail to induce apoptosis, resulting in primordial oocyte survival.

**Figure 1.**
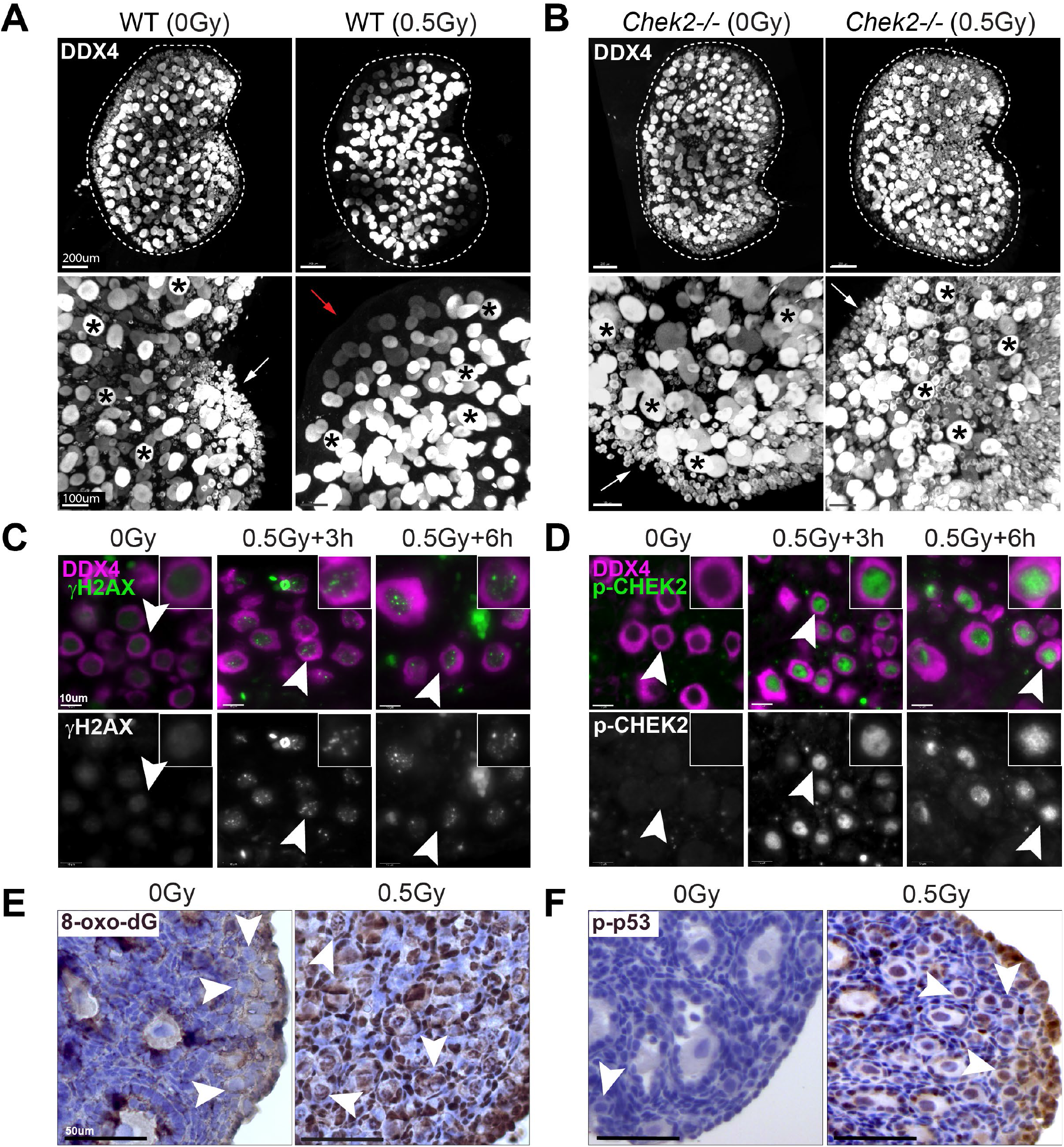
IR-induced DNA damage activates CHEK2-dependent signaling leading to primordial oocyte loss. IR dose of 0.5 Gy eliminates primordial oocytes (arrows) in wildtype (**A**) but not in *Chek2*-deficient ovaries (**B**). Large growing oocytes (asterisks) are not eliminated by IR. Red arrow indicates the area depleted of primordial oocytes. Shown are 3D renderings of whole ovaries (top) and magnified view (bottom) immunostained with oocyte marker DDX4. IR induces DNA damage in oocytes as evident by γH2AX staining (**C**) and leads to phosphorylation of CHEK2 (p-CHEK2) (**D**). IR also induces oxidative DNA damage in ovaries as evident by presence of the 8-oxo-2′-deoxyguanosine (8-oxo-dG) staining (**E**). IR leads to phosphorylation and activation of p53 (p-p53) in oocytes and other ovarian cells (**F**). Arrowheads indicate primordial oocytes.

### Transcriptional response to radiation-induced DNA damage in ovaries is CHEK2 dependent

In addition to p53 and TAp63, CHEK2 also phosphorylates and activates other targets including CDC25, NEK6, FOXM1, BRCA1, and BRCA2 during DDR (29–32). To identify proteins involved in the ovarian DDR signaling downstream of CHEK2 we performed transcriptional profiling of ovaries exposed to an oocyte-lethal dose of IR in wildtype and CHEK2 deficient females. We used juvenile ovaries for analysis because they are enriched for primordial oocytes that are extremely sensitive to DNA damage. Most primordial oocytes are depleted within 24 hours after IR (33). At 6 hours post-IR, primordial oocytes positive for DDR markers were still present in the ovary, suggesting active and ongoing DDR (**Figure 1C**). We hypothesized that analysis of differentially expressed genes in wildtype (radiation sensitive) and *Chek2-/-* ovaries (radiation resistant) would identify Radiation-Responsive Genes (RRGs) and signaling pathways downstream of CHEK2; those most likely contributing to primordial oocyte elimination (**Figure 2A**). RRGs differentially expressed in both wildtype and *Chek2-/-* ovaries will represent CHEK2-independent response pathways which we consider less likely to contribute to primordial oocyte elimination. We exposed one-week-old wildtype and *Chek2-/-* females to 0.5 Gy IR or sham treatment (N=6 for each group) and collected ovaries at 6 hours post-IR for RNA extraction and subsequent bulk RNA sequencing. Differential gene expression analysis in wildtype ovaries identified 83 RRGs with ≥2-fold change in expression and FDR≤0.05 (**Supplementary Data 1**). 77 genes were upregulated and 6 downregulated (**Figure 2B, C**; **Table S1**). 70 RRGs were protein coding genes, 7 were lncRNAs, and 6 were unclassified or pseudogenes. Surprisingly, overall global gene expression was not significantly altered by IR in *Chek2-/-* ovaries (**Figure 2B, C**) and only 3 genes showed a significant change (≥2-fold change and FDR≤0.05) (**Table S1; Supplementary Data 1**). Many RRGs were previously reported to participate in the p53 signaling pathway, apoptosis, or cell cycle (*Bbc3, Ccng1, Cdkn1a* (p21), *Trp73, Mdm2, Pmaip1, Tp53inp1, Lhx3, Eda2r, Nox1)* and were upregulated in response to IR in a CHEK2 dependent manner (**Figure 2D, E)**. These results confirm that CHEK2 is the major regulator of ovarian response to oocyte-lethal dose of IR. In addition to known cell cycle arrest and proapoptotic genes, we detected CHEK2-dependent upregulation of genes with unknown roles in the ovary such as *Cbr2, Ankrd65, Fermt1* and *Fermt3* (**Figure 2D, E**). These genes represent proteins with a broad spectrum of functions. For example, *Cbr2* or carbonyl reductase 2 (LogFC 5.07 FDR= 2.91E-05) is implicated in detoxification or inactivation of the reactive carbonyl species (RCS). Since RCS are induced by IR, CBR2 may play a beneficial role during IR response, although it may not be required for primordial oocyte survival in the absence of CHEK2. Interestingly, a gene encoding a protein structurally related to CBR2, dicarbonyl and L-xylulose reductase (DCXR) (34), was also induced by IR in the wildtype ovary (DCXR LogFC 1.5 FDR=0.003) (**Figure 2E, Table S1**). The function of *Ankrd65*, an ankyrin repeat domain 65 (LogFC 6.7 FDR=1.22E-18), is unknown. Ankyrin repeats mediate protein-protein interactions and many proteins with ankyrin repeats have been implicated in DNA damage response (35–37). *Fermt1* (LogFC 4.5 FDR=9.82E-06) and *Fermt3* (LogFC 3.6 FDR=4.15E-10), FERM domain containing kindlin 1 and 3 (also known as kindlin-1 and -3), are involved in the organization of focal adhesions that mediate cell-to-cell communication (38,39). Interestingly, FERMT1 plays a role in oxidative stress response and its deficiency results in increased sensitivity to oxidative stress (40). These results indicate that IR leads to CHEK2-dependent upregulation of known and novel genes in the ovary, which may play a role in DDR and regulation of primordial oocyte survival.

**Figure 2.**
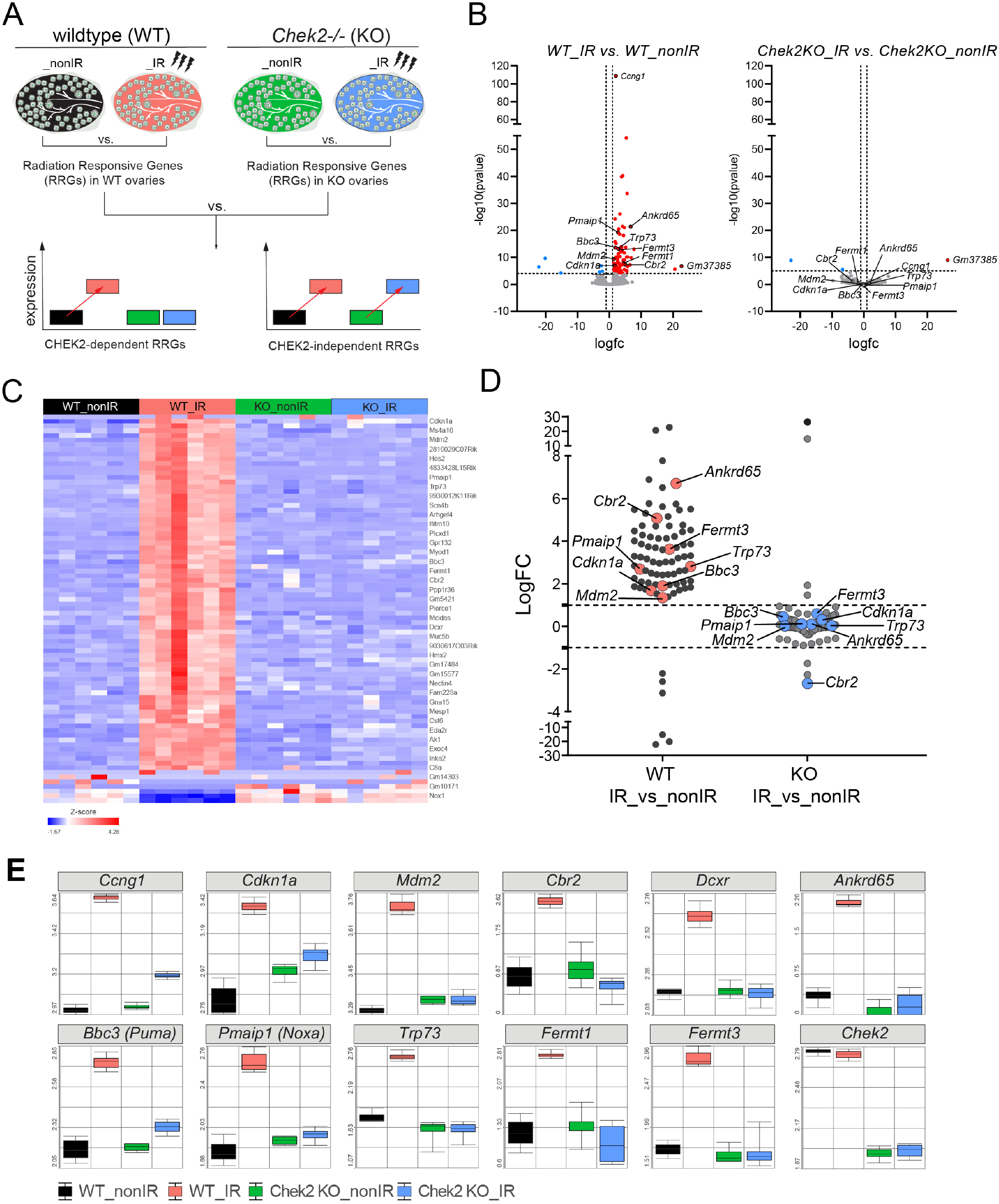
Radiation response in ovaries is abolished in the absence of CHEK2. **A)** Schematic representation of the experimental design. **B**) Volcano plot showing significant Radiation-Responsive Genes (RRGs) at FDR<0.05 in wildtype and *Chek2-/-* ovaries. Up-regulated LogFC≥1 (red), non-significant (grey), and down-regulated LogFC≤−1 (blue). **C**) Heatmap showing expression of 83 RRGs across samples. **D**) Comparison of gene expression changes (LogFC) induced by IR in wildtype vs *Chek2-/-* ovaries shows lack of response in the absence of CHEK2. **E**) Box plots showing expression of genes representing known DDR markers and newly identified genes in response to IR (N=6 samples per group).

### Radiation activates pathways related to apoptosis, interferon-mediated response, NFkB signaling, and changes in apical junctions in the ovary

To further identify the cellular and molecular response to IR in ovaries, we conducted a functional enrichment analysis for RRGs. The significantly enriched terms for Biological Process and KEGG pathways, as identified by g:Profiler (41), were associated with apoptosis and p53 activity (**Table S2**). These were driven by well-known genes such as *Pmaip1, Bbc3, Cdkn1a, Mdm2*, and *Trp73*. However, it’s important to note that radiation responses are dynamic and can vary among different cell types. As such, gene enrichment analysis based solely on differentially expressed genes with a ≥2-fold change might overlook significant effects on the activity of pathways involved in active responses. To address this, we conducted a Gene Set Enrichment Analysis (GSEA), which considers genes that are part of a differentially expressed set but may not individually reach statistical significance (42,43). Cumulative small changes in the expression of multiple genes belonging to the same gene set (pathway), but in a coordinated manner, could reveal other pathways involved in the radiation response. We performed GSEA using the Molecular Signatures Database (MSigDB) and the Hallmark gene set collection, which includes curated and well-defined biological processes (44). GSEA revealed 12 gene sets with significant changes (FDR≤0.05) in wildtype irradiated ovaries, including the activation of the p53-signaling and apoptosis pathways (**Figure 3** and **Table S3**). Furthermore, we detected activation of interferon alpha (INF-α) and gamma (INF-γ) responses, as well as TNFα signaling via NF-κB. Activation of these pathways may suggest an inflammatory response to radiation in the ovary (**Figure 3** and **Table S3**). Inflammation is a recognized response to radiation-induced tissue damage and has been linked to anti-apoptotic signaling, cell death, and fibrosis (45– 47). The inflammatory response in the ovary could indicate an active anti-apoptotic signaling in some cell types or a response to dying cells. In addition, GSEA analysis revealed an enrichment of pathways associated with apical junctions. This could imply alterations in cell-to-cell signaling during DDR or a reorganization of cell junctions because of cell death. Among the RRGs, 8 genes encode proteins associated with cell-cell junctions (*Trp73, Nectin4, Fermt1, Fermt3, Pkp3, Tjp3, Nox1, Hmcn2*) and 23 associated with the plasma membrane (*Plch2, Ifitm10, Ano3, Eda2r, Gpr132, Islr2, Scn4b, Slc6a3, Nectin4, Crhr1, Gramd2a, Pkp3, Tjp3, Fermt1, Ak1, Baiap3, Nox1, Itgb7, C8a, Hmcn2, Ms4a10, Dcxr, Nlrp6*). The implications of these changes at cell-cell junctions will require further investigation as they may indicate coordinated signaling between pre-granulosa cells and oocytes within the follicle or extrinsic signaling from other cell types in the ovary.

**Figure 3.**
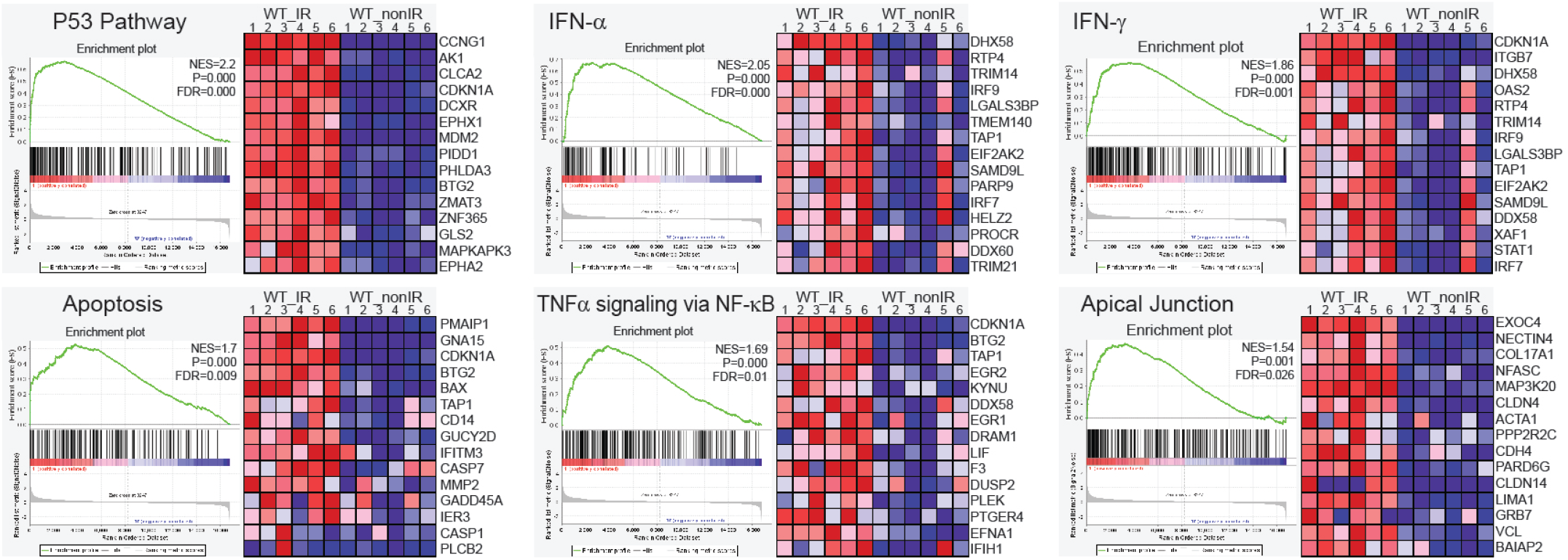
Gene Set Enrichment Analysis (GSEA) in HALLMARK datasets. The p53, apoptosis, inflammatory and apical junctions gene sets were significantly activated in irradiated ovaries. The plots show the leading edge (most significant genes) as vertical bars accumulated below the peak of the green enrichment score (ES) line and on the right as heatmaps. NES-normalized ES, P-nominal p value, FDR q value. The significance criteria were nominal P-value <0.05 and FDR q-value≤0.05.

### Ovarian radiation-response genes are enriched for p53 and p63 targets

In response to DNA damage, the CHEK2 kinase activates several effector proteins, including two transcription factors: p53 and p63 (31). These factors are known to induce apoptosis (32,48–51). We conducted a gene enrichment analysis using the g:Profiler TRANSFAC database to determine whether RRGs identified in the ovary are regulated by p53, p63, or other transcription factors (TFs). Our analysis revealed that ovary RRGs were enriched for p53 binding motifs (55 target genes), as well as for p63 (40), and p73 (14). However, these largely overlapped with p53 target genes (**Figure 4A**). These findings suggest that the CHEK2-dependent response to radiation in the ovary is predominantly mediated by p53 and TAp63. Interestingly, expression of *Trp73*, the third member of the family, was induced in the ovary after radiation in a CHEK2-dependent manner (**Figure 2C, E**). To identify p53 and TAp63 gene targets experimentally validated by Chromatin Immunoprecipitation (ChIP) and sequencing experiments (ChIP-seq), we performed ChIP-X Enrichment Analysis (ChEA) (52). We used the ChEA3 database, which aggregates previously annotated TF targets from multiple ChIP-seq experiments, including the ReMap dataset that includes mouse, human, and fly tissues (53). This approach identified p63 as the top-ranking TF and p53 as the fourth (**Figure 4B, C**). The top 10 TFs associated with ovary-RRGs include other TFs previously linked to the DNA damage response, such as CBX (54–56), TCF3 (57,58), SNAI2 (59,60) and THAP1 (61,62). Interestingly, novel ovary-RRGs such as *Cbr2, Ankrd65, Fermt1* and *Fermt3*, were all confirmed to be regulated by either p53 or p63 (**Figure 4A, B**). As the ReMap dataset does not include ovarian tissue, we performed RT-qPCR analysis on irradiated ovaries from wildtype, *Chek2-/-, Trp63A/A* (phosphomutant S621>A (12)), and double mutants lacking both active TAp63 and p53 (*Trp63A/A; Trp53-/-*) to test whether these novel ovary-RRGs are activated by p53 or TAp63. The upregulation of *Cdkn1a*, a known p53-specific target involved in cell cycle arrest, was observed in wildtype and TAp63 phosphomutant ovaries, but not in ovaries lacking active p53 or CHEK2. In contrast, the upregulation of *Cbr2, Ankrd65, Fermt1*, and *Fermt3* was abolished in the absence of active TAp63 or CHEK2. These results confirmed the CHEK2-dependent upregulation of these ovary-RRGs and demonstrated that *Cbr2, Ankrd65, Fermt1* and *Fermt3* are predominantly induced by TAp63 in the ovary (**Figure 4C**). As TAp63 is exclusively expressed in oocytes, we predicted that these genes will be upregulated in oocytes. To test this, we performed RNA *in situ* hybridization (RNA-ISH) using the RNAscope system. We observed a strong signal for all three genes near the periphery of the ovary, where PFs reside, exclusively in irradiated oocytes (**Figure 4D**). Together, our results indicate that CHEK2 mediates the response to radiation-induced damage mainly via TAp63 in oocytes but also suggests a role for p53 in ovarian DDR.

**Figure 4.**
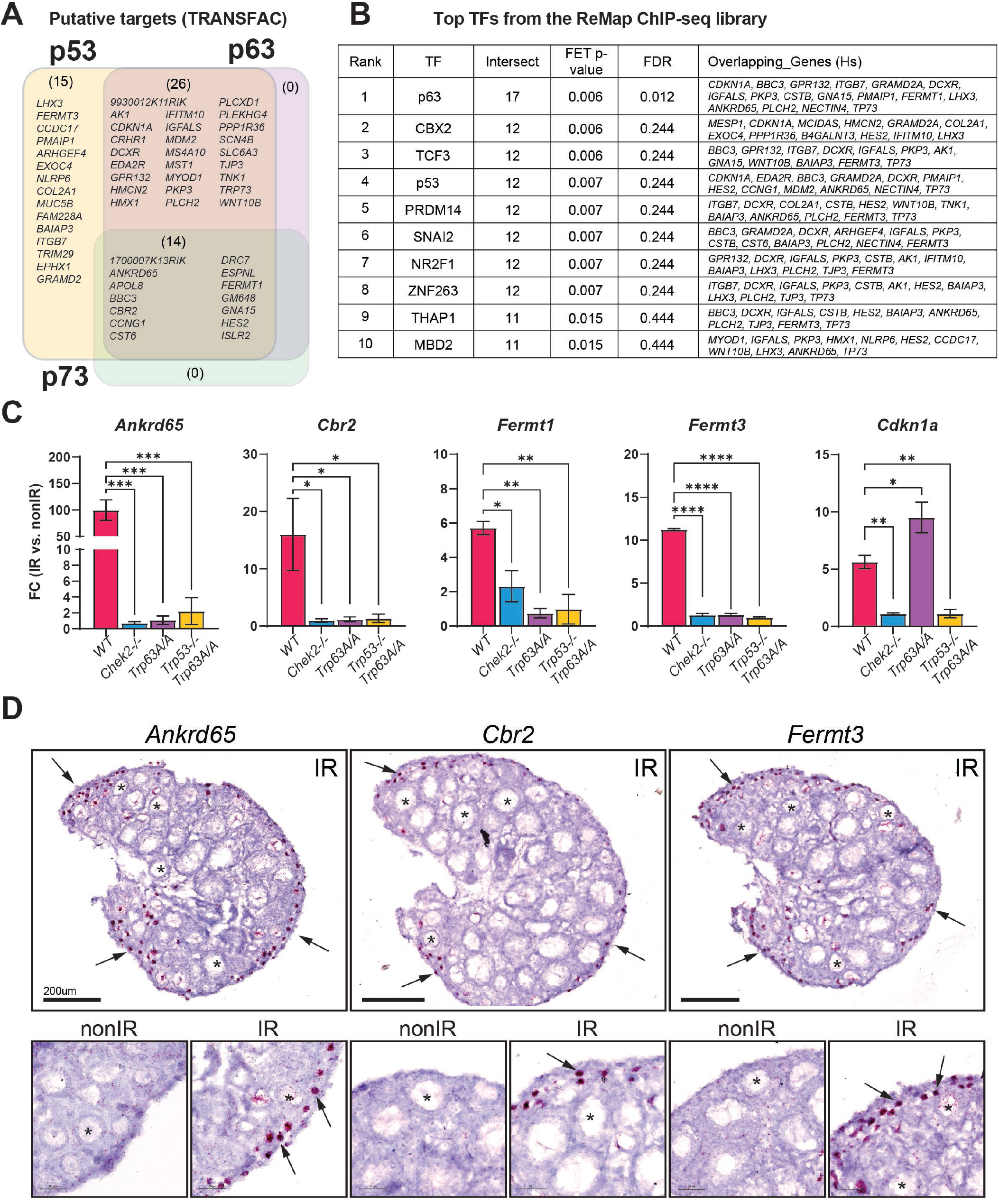
Radiation induces expression of TAp63 and p53 target genes. **A**) Venn diagram showing putative targets of p53, p63 and p73 transcription factors (TF) among RRGs based on TRANSFAC database. **B**) Top TFs predicted to regulate RRG expression based on validated ReMap ChIP-seq datasets. **C**) RT-qPCR analysis of RRGs in mutants lacking active CHEK2, TAp63 and p53. **D**) RNAscope validation of oocyte-specific upregulation of novel RRGs in the ovary. Red signal indicates presence of the transcript. Arrows indicate oocytes.

### Radiation Elicits a Stronger Response in Oocytes Compared to Somatic Cells

The majority of ovary-RRGs identified by bulk RNA-seq are expressed in various cell types and tissues, with some never reported to be expressed in untreated ovaries (**Figure S1**). This raises the question of whether ovary-RRGs are induced in a specific cell type or if bulk RNA-seq can detect responses in primordial oocytes. Although CHEK2 is expressed in all cell types in the ovary, primordial oocytes are the most sensitive to IR and undergo apoptosis within 24 hours post-IR (33). This contrasts with most somatic cells in the ovary, including pre-granulosa cells, which survive and persist longer, even in the absence of oocytes. To dissect how CHEK2 regulates cell type-specific responses to IR-induced damage, we conducted single-cell RNA sequencing (scRNA-seq) in wildtype and *Chek2-/-* ovaries exposed to the same IR regimen as in bulk experiments (6 hours post-IR, 0.5 Gy vs. sham). We identified clusters corresponding to different cell types based on well-established markers, including *Dppa3, Sycp3, Dazl, Zp3, Gdf9* for oocytes, and *Inha, Inhbb, Amh, Amhr2* for granulosa cells (**Figure 5A-C** and **Figure S2A**). Overall, 11 clusters were identified, representing 9 cell types: oocytes, granulosa, fibroblasts, endothelial, epithelial, erythrocytes, macrophages, pericytes, and perivascular cells (**Figure 5B**). Oocytes represented 1-2% of all cells in the ovary, compared to approximately 38% of granulosa cells, and 50% of fibroblasts (**Figure 5D**). To determine whether ovary-RRGs from bulk analysis were induced in oocytes or other cell types, we analyzed their expression in individual clusters. We found that the majority of ovary-RRGs were predominantly induced in irradiated oocytes (cluster 9) (**Figure 5E**). Moreover, their induction was not observed in *Chek2-/-* cells. Next, we investigated IR-induced changes in gene expression across clusters and cell types. We found that the oocytes have the highest number of differentially expressed RRGs compared to other cell types (**Figure 6A** and **Figure S2B-D**). In the oocyte cluster, 86 genes were differentially expressed at FC ≥1.5 and FDR ≤0.05. Among the genes were *Fermt1, Fermt3*, and *Cbr2*, which were previously identified by bulk analysis. The second highest upregulation of gene expression was observed in epithelial cells in ovarian surface epithelium (20 genes) (**Figure 6A**). Epithelium showed strong signature of interferon-induced immune response (*Isg15, Rsad2, Stat1, Tnfrsf12a*) and interferon-inducible genes (*Ifit1, Ifit3, Ifitm3, Iigp1, Irgm1*) (63,64) (**Supplementary Data 2**). Granulosa cells, the second most abundant cell type in the ovary, displayed changes in the expression of 11 genes, including *Cdkn1a* and *Ccng1*, indicating induction of cell cycle arrest. As in bulk RNA-seq, very few DEGs were identified in *Chek2-/-* cell clusters (**Supplementary Data 2**). These results indicate that IR-induced damage in the ovary elicits the strongest response in oocytes. This is supported by cellular changes observed in the ovary after IR, such as a high level of DNA damage marker γ-H2AX in oocytes compared to somatic cells (**Figure 1B**), and persistence of pre-granulosa cells after the loss of primordial oocytes (**Figure S3**). Next, we compared the overlap between ovary-RRGs from bulk analysis and RRGs from each cell type from wildtype irradiated and non-irradiated ovaries (FC≥1.5 FDR≤0.05). The largest overlap was with the RRGs in the oocyte cluster (oocyte-RRGs) and included *Cbr2, Dcxr, Fermt1, Fermt3, Ifitm10, Mdm2, Cdkn1a, Gm648, Hmcn2, Bbc3, 9930012K11Rik, Cst6*, and *1700007K13Rik* (**Figure 6A**). *Uba52, Cdkn1a*, and *Bax*— known DDR, cell cycle arrest, and apoptosis factors—were differentially expressed in multiple cell types. To assess DDR response patterns in gene expression across different cell types, we visualized the expression of core DDR genes and selected RRGs from this study using radar charts (**Figure 6B**). Known DDR genes such as *Cdkn1a, Bbc3 Ccng1*, and *Mdm2* were induced in multiple cell types, while *Cbr2, Fermt1, Fermt3, Ankrd65, Trp73*, and a few other RRGs were exclusively induced in oocytes (**Figure 6B** and **Figure S4**). Interestingly, *Mdm2*—a negative regulator of p53—showed the highest induction and expression in irradiated oocytes compared to other cell types. This may suggest a unique mechanism limiting p53 activity in primordial oocytes during DDR (**Figure 6B**). Among oocyte-RRGs that were not identified by bulk analysis were *Perp, Ddit4, Fos*, and *Phlda3*, which have all been previously implicated in DDR and p53 pathways (65–67). To uncover the differences in how individual cell types respond to IR-induced damage, we conducted GSEA utilizing hallmark gene sets and pathways (**Figure 6C**). Given the lack of significant response in *Chek2-/-* ovaries, our analysis focused on wildtype ovaries. We observed clear differences in pathway activation between cell types. The strongest activation of apoptosis was detected in oocytes, erythrocytes, and perivascular cells. G2M checkpoint activation was detected in fibroblasts, epithelial, and endothelial cells. The p53 pathway was activated in oocytes, granulosa cells, fibroblasts, erythrocytes, and perivascular cells. Oocytes and epithelial cells showed induction of the c-MYC signaling, which is involved in p53-induced apoptosis (68).The INF-α, INF-γ, and TNFα signaling via NFkB— previously identified in bulk RNA-seq analysis—were activated in granulosa and epithelial cells. In summary, the transcriptional profiling reveals that IR-induced damage is more harmful to primordial oocytes and results in a stronger response compared to other cell types. Moreover, analysis at the single-cell level reveals that IR-induced damage results in the activation of different response pathways in different cell types.

**Figure 5.**
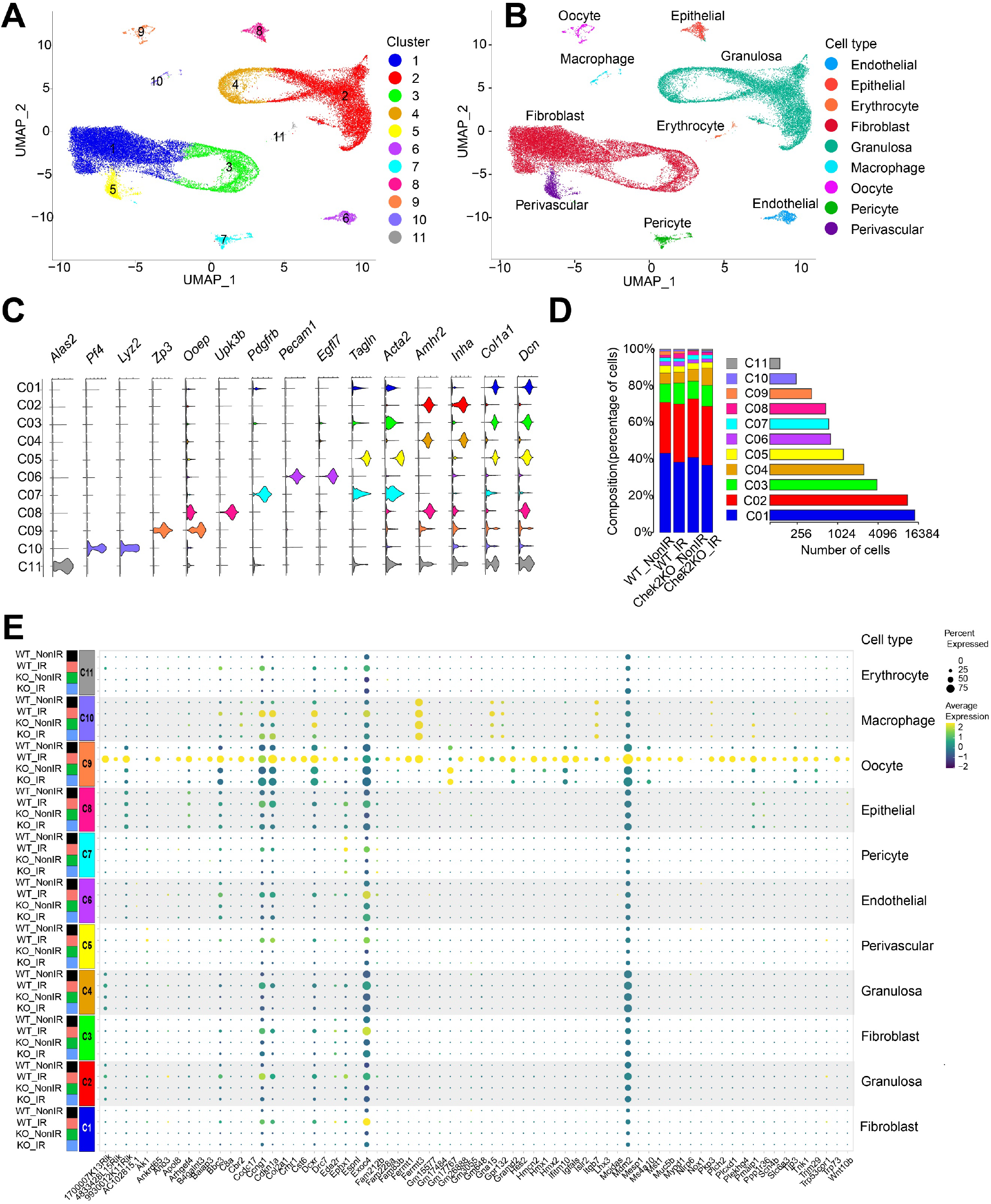
Radiation Responsive Genes are predominantly induced in oocytes. UMAP plots of all cells from wildtype and *Chek2-/-* ovaries with and without radiation identified 11 clusters (**A**) representing nine cell-types (**B**). Clusters were identified using cell-type-specific markers (**C**). **D**) Distribution of cells among clusters. **E**) A dot plot showing the relative expression of all RRGs identified by bulk RNAseq across all clusters.

**Figure 6.**
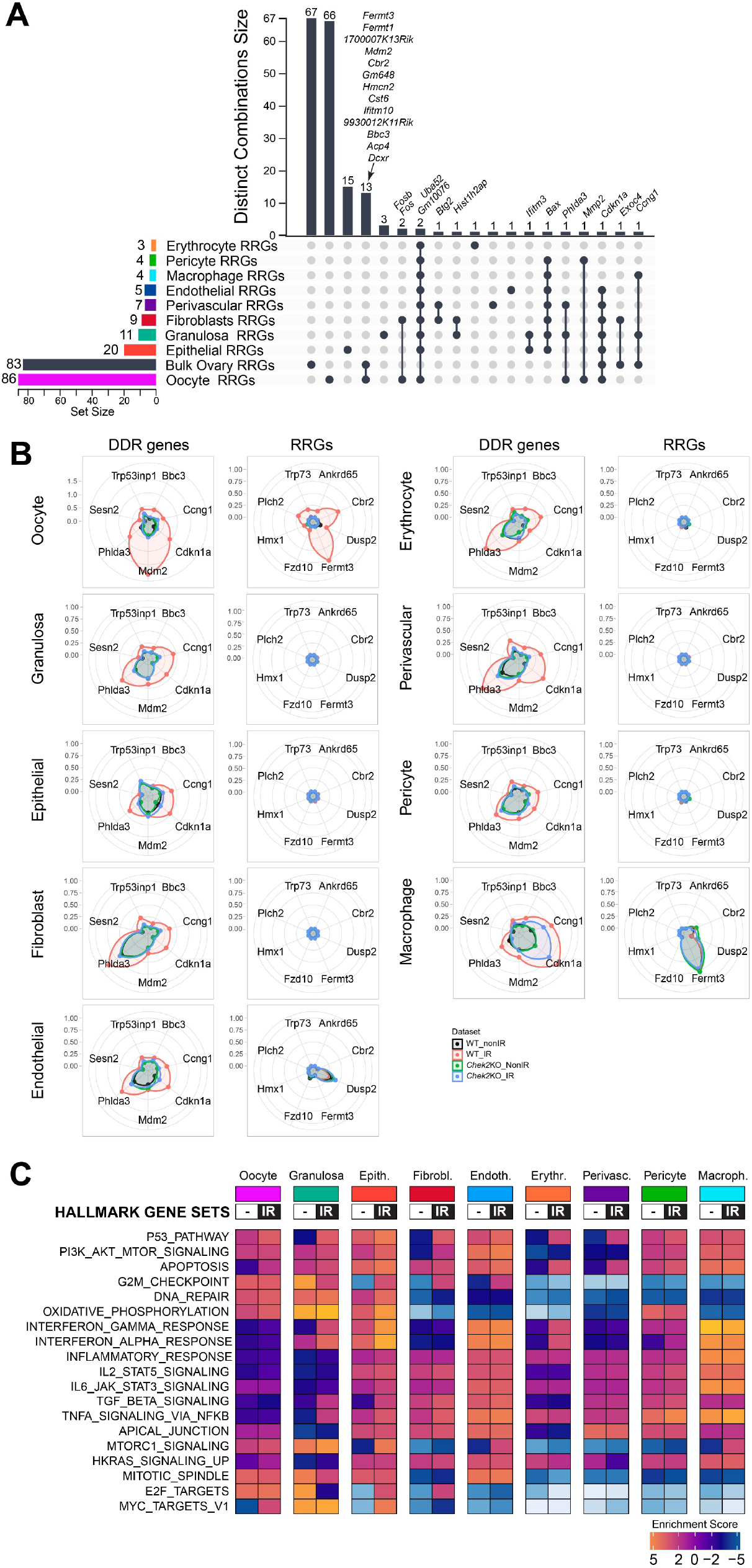
Radiation response varies between different cell types in the ovary. **A**) UpSet plot showing DEGs specific to a cell-type and DEGs shared between cell-types (single filled circle and filled circles connected with vertical lines, respectively). Vertical bar plot indicates unique or overlapping DEGs in clusters. Horizontal bar plot indicates the number of RRGs per cell-type. **B**) Radar charts comparing upregulation of known DDR genes and RRGs in different cell types in wildtype and *Chek2-/-* ovaries. Concentric rings represent log2FC. **C**) Heatmap showing different hallmark gene sets activated in specific cell-types after irradiation.

### Radiation induces unique cellular responses in oocytes

To enhance our understanding of the oocyte-specific response to IR-induced damage, we re-clustered both irradiated and non-irradiated oocytes from wildtype and *Chek2-/-* females into six distinct subclusters. Interestingly, we identified a unique group, subcluster 5, consisting exclusively of irradiated oocytes from the wildtype group, while the remaining five subclusters included oocytes from all four groups (**Figure 7A**). 44% of all irradiated wildtype oocytes were in subcluster 5, suggesting that the remaining irradiated oocytes in other subclusters may be at an earlier stage of DDR or from different follicle types. Subcluster 5 was distinguishable for the differential expression of 147 genes (FC ≥1.5 FDR≤0.05) (**Supplementary Data 3**). Among them, 15 genes previously identified as RRGs in bulk ovary analysis (ovary-RRGs), *Fermt1, Fermt3, Cbr2, 1700007K13Rik, Mdm2, 9930012K11Rik, Cdkn1a, Cst6, Ifitm10, Acp4, Dcxr, Hmcn2, Bbc3, Gpr132*, and *Gm648* (**Figure 7B, Figure S5**). We further examined the expression of all ovary-RRGs from the bulk analysis across the oocyte subclusters and observed that the majority were specifically upregulated in subcluster 5 (**Figure 7C**). The fact that subcluster 5 oocytes represent 44% of all oocytes in the wildtype irradiated sample indicates that they largely contributed to RRG signature in the ovary. However, many ovary-RRGs did not reach statistical significance in single-cell and subcluster analysis, potentially due to oocyte-to-oocyte variability in expression levels. The analysis of subcluster 5 identified additional oocyte-RRGs. Some are linked to p53 and apoptosis such as *Ddit4* (66), *Perp* (69) and *Pycard* (70), others function as cell surface receptors such as *Lama5* (71), and *Mcam* (72) (**Figure 7B, D** and **Supplementary Data 3**). RRGs from both bulk and subcluster analyses demonstrated CHEK2-dependent upregulation in the oocyte cluster (**Figure S6**). GSEA revealed a strong enrichment for pathways related to p53, DNA repair, and oxidative phosphorylation (**Figure 7E**). Enrichment for mTORC1, E2F, MYC targets, apical junctions, G2M checkpoint, and apoptosis was also observed in subcluster 5. While p53 and apoptotic signaling were expected in response to IR, the upregulation of pathways related to cell junctions detected by both bulk and single-cell approaches is noteworthy. Cell-to-cell communication through gap junctions can propagate an IR-damage response between cells via bystander effect, and the desmosomal protein PERP—upregulated in oocytes after IR (**Figure 7D**)— is a known apoptotic factor (65,73). To expand on our findings from the bulk analysis, we performed TF enrichment analysis for subcluster 5 RRGs using ChIP-seq data collected from the ReMap database, and again identified p63 as the top-ranking TF, followed by KLF9 and p53 (**Figure 7F**). Two other transcriptional regulators already identified in bulk ovary analysis (**Figure 4B**), SNAI2 and THAP1, were ranked higher for oocyte-RRGs (**Figure 7F**). Interestingly, p63 and THAP1 are predominantly expressed in oocytes compared to other ovarian cells (**Figure S7**), suggesting that they may propagate DDR specific to oocytes. p53 and KLF9 are expressed in all cell types while SNAI2 is highly expressed in fibroblasts (**Figure S7**). Overall, 32 DEGs in subcluster 5 were previously experimentally validated as targets for p63, 30 for p53, and 19 shared by both factors (**Figure 7G**). In summary, *de novo* subclustering of oocytes from control and irradiated wildtype and *Chek2-/-* ovaries identified oocytes undergoing active DDR. These oocytes show strong activation of p53 pathway, DNA repair, and oxidative phosphorylation. Interestingly, activation of the apical junction pathway and upregulation of plasma membrane/cell surface proteins in oocytes in subcluster 5, suggests an intriguing possibility of coordinated DDR signaling at the interface between primordial oocytes and pre-granulosa cells. Moreover, these results indicate that, although TAp63 primarily mediates DDR and apoptosis in oocytes, the strong signature of p53-related signaling suggests an active role for p53 in the oocyte’s response to DNA damage. Further studies are needed to determine which DDR processes in oocytes are regulated by p53, and to which extent DDR in pre-granulosa cells affects oocyte survival or death.

**Figure 7.**
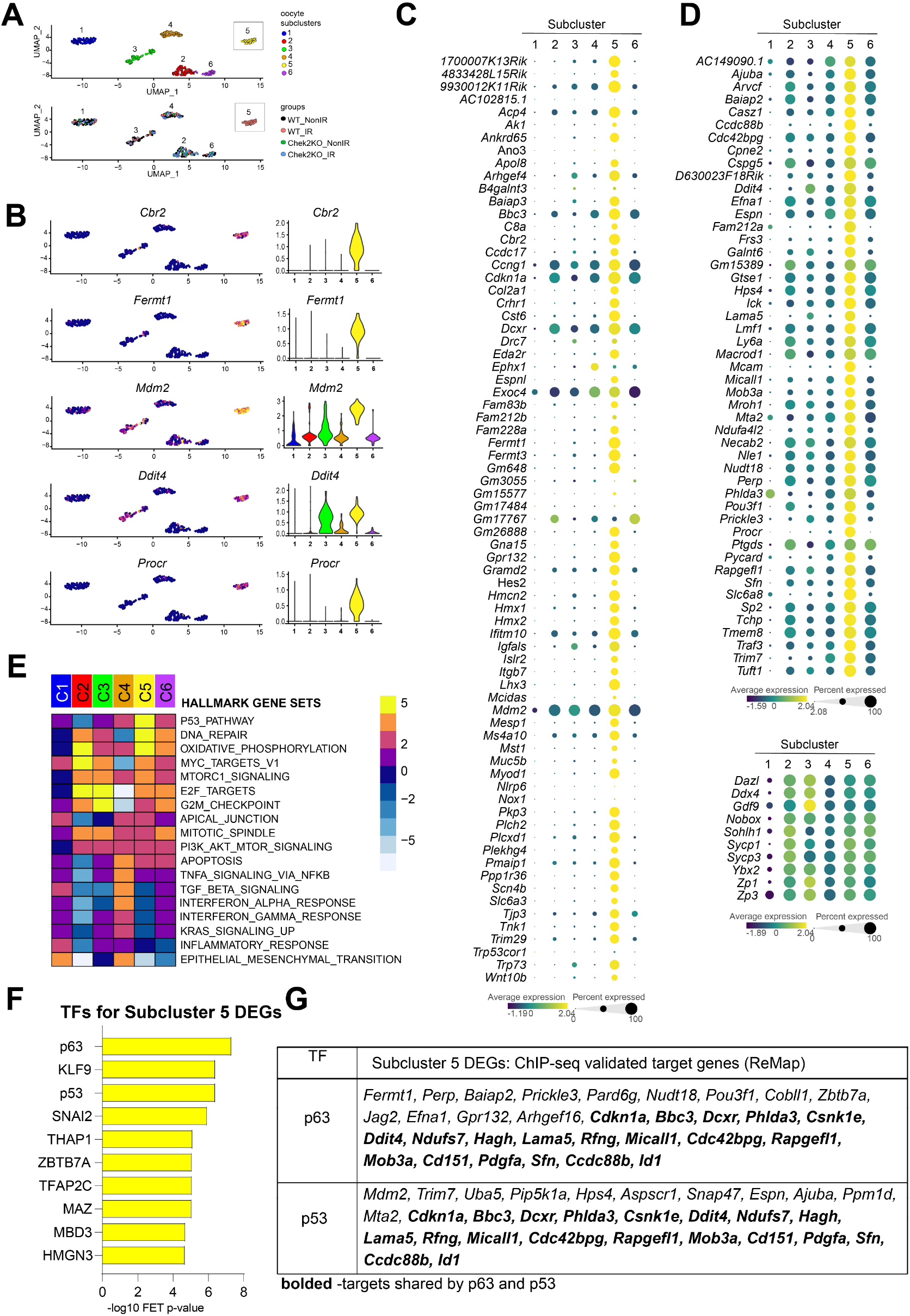
Oocytes display unique and diverse transcriptional changes in response to radiation. **A**) UMAP plots of oocyte subclusters. Subcluster 5 is comprised of solely wildtype irradiated oocytes. **B**) Gene expression levels of representative RRGs overlaid on UMAP plots and shown by violin plots. **C**) Dot plot showing subcluster-specific expression of ovary-RRGs from bulk analysis. **D**) Dot plot showing subcluster-specific expression of additional oocyte-RRGs identified by subcluster analysis and oocyte marker genes (bottom). **E**) Heatmap showing subcluster-specific gene set enrichment for different Hallmark gene sets. **F**) TF enrichment analysis using ChEA3 ReMap dataset. **G**) DEGs in subcluster 5 previously validated as p63 and p53 targets.

## Discussion

Females are born with a finite supply of immature PF, which are expected to last throughout their reproductive lifespan until menopause. As we increasingly recognize that various exogenous and endogenous sources can cause DNA damage in primordial oocytes, leading to their accelerated depletion, it becomes critical to identify the factors that regulate the oocyte’s response to DNA damage and its survival. This is important, because loss of immature follicles results in loss of fertility and hormonal dysregulation in females. In this study, we employed both bulk and single-cell transcriptomics to uncover the molecular mechanisms that lead to the elimination of primordial follicles in response to DNA damage caused by ionizing radiation. Our approach revealed that the response to DNA damage induced in ovaries, by a relatively low but oocyte lethal dose of radiation, is solely dependent on CHEK2 downstream signaling as the transcriptional response was almost entirely abolished in its absence. Radiation-induced DNA damage disproportionately affected oocytes and induced the strongest response at the transcriptional level driven by two CHEK2 targets: TAp63 and p53. We further demonstrate that the ovarian response to radiation involves the activation of interferon and inflammatory pathways in the ovarian soma and signaling at the cell-cell junctions likely between the oocyte and pre-granulosa cells. Our findings reveal novel genes and unique responses in oocytes DDR, which may help explain their sensitivity to various genotoxic insults.

Single-cell composition of one-week-old ovaries based on transcriptional cell clustering indicates that they are comprised of 9 major cell types, with fibroblasts (or fibroblast-like) and granulosa cells being the two most abundant and constituting almost 90% of all cells. Oocytes make up less than 2%, while the remaining cell types include endothelial, epithelial, erythrocytes, macrophages, pericytes, and perivascular cells. Following IR, oocytes exhibit the highest number of differentially expressed genes. This suggests that primordial oocytes are exquisitely sensitive to DNA damage and harbor a unique DDR. Somatic cells in the G2/M phase of the cell cycle are most sensitive to DNA damage presumably due to the limited time for repair before chromosome segregation and subsequent cell division. Therefore, it may not be surprising that primordial oocytes which are suspended in the dictyate stage of meiotic prophase I, similar to G2 phase, are highly sensitive to DNA damage. However, despite remaining arrested for many months or years—which would give them enough time to repair damage before resuming meiosis, they seem to default to apoptosis rather than repair.

The examination of changes in gene expression indicates that oocytes exhibit the strongest response to DNA damage in the ovary. Radiation treatment led to upregulation of multiple known DDR/apoptotic genes in oocytes including *Cdkn1a, Bbc3, Ddit4, Perp* and *Phlda3*. This is in agreement with a previous study in purified primordial oocytes which reported an enhanced expression of pro-apoptotic genes (74). However, we also identified other upregulated genes in irradiated oocytes that point to additional response pathways that were not previously implicated in the DDR in ovaries. For example, upregulation of *Cbr2* and *Dcxr—*which belong to a family of carbonyl oxidoreductases—reveals a potential mechanism of detoxification in response to oxidative damage specifically in oocytes (34). These proteins can reduce endogenous and exogenous carbonyl compounds including those derived from lipid peroxidation of mitochondrial membranes and can provide protection against reactive oxygen species (ROS). CBR2 is predominantly expressed in lungs and is reported to localize to mitochondria but it’s exact function remains unknown (75). *Cbr2-/-* females seem to be fertile, as are *Chek2-/-*, therefore more in-depth studies are needed to determine the role of CBR2 in ovarian DDR and follicle survival (76). DCXR is a multifunctional protein involved in sugar metabolism, carbonyl detoxification, cell adhesion, and male fertility (34). *C. elegans* ortholog of DCXR (DHS-21) plays a role in detoxification of carbonyl compounds, and *dhs-21* mutants have shorter lifespans presumably due to decreased defense against oxidative damage (77). *Dcxr*-deficient mice die before wean age, thus the role of DXCR in ovaries and female fertility was not assessed (76). However, DCXR was detected in human oocytes by Human Protein Atlas (78). This suggests that CBR2 and DCXR may protect oocytes from oxidative damage induced by IR and endogenous sources, however they may not be required for oocyte survival in the absence of pro-apoptotic signaling. Although a *Cbr2* ortholog was not found in humans, it is possible that DCXR alone fulfills this function in human oocytes. Intriguingly, DCXR is also implicated in cell adhesion and has been shown to colocalize with E-cadherin at cell-cell junctions (34,79). Importantly, *Dcxr* is not the only gene associated with cell-cell junctions induced by IR in oocytes, suggesting a potential role for communication between oocytes and pre-granulosa cells in the regulation of apoptosis. *Fermt1* and *Fermt3*, also known as kindlin-1 and 3, are best known for their role in cell-extracellular matrix adhesion and integrin activation (38,39). The exact mechanism by which integrin-mediated adhesion may regulate oocyte survival remains unknown. Nevertheless, it has been demonstrated that the activation of integrins and cell adhesion can regulate cell survival in response to DNA damage through the modulation of p53 and p73/c-Abl, thereby potentially linking kindlins and apoptosis in oocytes (80,81). Furthermore, we found that IR causes an upregulation of the cell-adhesion gene *Perp* in oocytes. PERP, a desmosomal protein, is a known target of p53 and p63 and is involved in apoptosis (69,82,83). The oocytes in PFs are surrounded by pre-granulosa cells, which are similarly quiescent and non-proliferative. They support oocyte quiescence and survival through direct contact and bidirectional communication across cell-cell junctions (84–86). These findings imply that there may be a coordinated response to radiation between primordial oocytes and the surrounding pre-granulosa cells.

In contrast to oocytes, few genes show differential expression following radiation in fibroblasts and granulosa cells (9 and 11 DEGs, respectively). The upregulation of *Cdkn1a, Ccng1*, and *Bax* in these cells suggests the activation of cell-cycle arrest to facilitate repair and survival, or apoptosis if the damage is irreparable. This implies that the IR dose used in this study, while lethal to primordial oocytes, is tolerated by these cell types. This is corroborated by previous studies where neither lower (0.1Gy) nor higher (1Gy) doses resulted in fibrosis or defects in granulosa cell proliferation which could arise after IR (87,88). Due to a lack of definitive markers to distinguish pre-granulosa cells in primordial follicles, we were unable to identify them in the granulosa cell cluster. Transcriptomic studies with high spatial resolution are needed to determine how pre-granulosa cells and primordial oocytes respond to radiation.

The ovarian surface epithelium showed the second highest number of differentially expressed genes (20 DEGs), exhibiting a strong interferon-induced immune response. This included genes such as *Isg15, Rsad2, Stat1, Tnfrsf12a*, and interferon-inducible genes *Ifit1, Ifit3, Ifitm3, Iigp1, Irgm1*. Most of our knowledge about the role of interferon signaling in response to IR-induced DNA damage comes from studies in cancers where it is often linked to cGAS/STING signaling (89–92). Interferon alpha was shown to induce apoptosis via the mitochondrial pathway and activation of BAX, which we see upregulated in epithelial cluster cells (93). ISG15, a ubiquitin-like protein that conjugates to various proteins in response to interferon, was previously shown to be upregulated in response to DNA damage and has been implicated in regulation of p53 and p63 (94,95). Interestingly, while TAp63 is exclusively expressed in oocytes, p53 and CHEK2 are ubiquitously expressed in all cells within the ovary. Therefore, it is possible that extrinsic signaling from other cells in the ovary, including the neighboring pre-granulosa and surface epithelium or fibroblasts in the stroma, may also play a role in regulating the fate of primordial oocytes either by direct cell-cell communication, or by stimulating changes in the microenvironment (96–98). Intriguingly, a previous study reported a strong interferon response in purified oocytes after IR (74). Our study detected INF-α, INF-γ, and inflammatory signatures in ovaries by bulk RNA-seq but not in oocytes using scRNA-seq. It is possible that the protocol used for oocyte purification resulted in contamination with epithelial cells or incomplete separation of primordial oocytes from pre-granulosa cells, which may be the actual source of the interferon signaling. Further studies are needed to determine the role of interferon/ISG15 signaling in primordial follicles or ovarian epithelium in response to DNA damage.

Although p53 is activated by DNA damage in primordial oocytes, and p53-regulated genes are upregulated in response to IR in oocytes, much less is known about the role of p53 in oocyte DDR compared to TAp63. Our previous work has shown that alkylating agents induce more DNA damage in oocytes, which leads to activation of p53-dependent apoptosis in oocytes, indicating that specialized mechanisms may regulate p53 activity in oocytes (12). A better understanding of the DDR responses driven by p53—that may be beneficial to oocyte’s survival but are made ineffective by excessive (or precautious) TAp63-driven apoptosis—will be critical for elucidating the mechanisms of ovarian aging, and ovariotoxic effects of therapeutic or environmental genotoxic exposures.

## Materials and Methods

### Animals

All procedures used in this study were approved by the IACUC at The Jackson Laboratory. C57BL/6J (#000664) and *Trp53*tm1Tyj/J (#002101) mice were obtained from The Jackson Laboratory. *Chek2*tm1b(EUCOMM)Hmgu mice were obtained from KOMP program at JAX. *Trp63*S621A mutant line was generated using CRISPR/Cas9 (12). For total body irradiation experiments, 7 to 9 day old females were irradiated using a Cesium-137 gamma irradiator. Females were exposed to sham or a single dose of 0.5 Gy administered at a rate of (∼170 Rad/min). Ovaries were collected at 3 or 6 hours after irradiation for RNA or protein extraction or were fixed in 4% PFA for immunostaining. For whole ovary immunostaining and imaging, ovaries were collected two weeks after irradiation following perfusion with PFA as previously described (99).

### Immunohistochemistry

Ovarian sections of 5μm thickness were prepared and immunostained using standard procedures. Primary antibodies used in this study were mouse anti-p63 (4A4, Biocare Medical, CM163A), rabbit anti-pCHEK2(T68) (Bioworld Technology, BS4043), rabbit anti-phospho-p53(S15) (Cell Signaling, 9284), rabbit anti-DDX4 (Abcam, ab13840), mouse anti-γ-H2AX (Millipore, 05-636). Mouse anti-DNA/RNA Damage antibody [15A3] (Abcam, ab62623) recognizing oxidative DNA and RNA damage (8-hydroxy-2’-deoxyguanosine, 8-hydroxyguanine and 8-hydroxyguanosine). The secondary antibodies used were Alexa Fluor (Invitrogen). Immunostaining for phospho-p53(S15) and oxidative damage was performed with Starr Trek reagent (Biocare Medical, STUHRP700H). Imaging was performed using a Leica DM550 microscope and LAS X software (Leica).

### Whole ovary immunostaining, optical clearing, and imaging

Immunostaining and optical clearing of whole ovaries was performed using CUBIC as described in (99). The primary antibody incubation was carried out at room temperature with gentle rocking for 2-4 days. The primary antibodies used were rabbit anti-DDX4 (Abcam, ab13840) and secondary antibodies were Alexa Fluor (Invitrogen). Whole ovaries were imaged using the Leica DIVE/4Tune multiphoton and LAS X software (Leica). 3D rendering was prepared with IMARIS software (Bitplane).

### In situ hybridization using RNAScope

Ovaries from 9-day-postpartum pups treated with sham or 0.5 Gy IR were fixed in 4% paraformaldehyde overnight at 4ºC, embedded by freezing in O.C.T. media (Tissue-Tek), and sectioned into 10 μm thickness. Sections were processed using the Manual RNAscope 2.5 HD RNAscope kit (Advanced Cell Diagnostics (ACD) #322350) as described in the manufacturer’s instructions. In brief, 10-μm ovarian sections were pretreated with protease before hybridization with target probes: *Ankrd65* (ACD #843881), *Cbr2* (ACD #842871), and *Fermt3* (ACD #562571) and control probes: *Ppib* (ACD #313911), *DapB* (ACD #310043). Sections were incubated with amplifier probes AMP1 through AMP5, and then Fast-RED A/B solution was applied to sections for chromogenic staining of probes. Positive staining was identified as red, punctuate dots in the cells. Sections were counterstained with Hematoxylin Gills I and 0.02% ammonia water was used for blueing. Images were acquired using a bright field microscope (Leica DM5500) or NanoZoomer (C-13210-01). Experiments were performed with at least two ovaries per IR condition with a minimum of three replicates.

### Quantitative RT-qPCR Analysis

Ovaries were dissected from female pups exposed to sham or 0.5 Gy radiation (N=3 per group). The ovaries were immediately frozen in liquid nitrogen and stored at -80°C until RNA extraction. Total RNA was extracted from pooled ovaries using a RNeasy Micro Kit (Qiagen, 74004) according to the manufacturer’s instructions, with an additional DNAse I treatment step to remove DNA contamination. A minimum of 500 ng of total RNA was reverse transcribed into cDNA using SuperScript IV reverse transcriptase (Invitrogen, 18091050) according to the manufacturer’s instructions. The cDNA samples were used as templates for quantitative real-time PCR (qPCR) using the ViiA7 Real Time PCR system (Life Technologies) and the Power Track SYBR Green PCR master mix (Applied Biosystems, A46109). The qPCR reactions were performed in triplicate. The primer sequences for the target genes and the reference gene *Gapdh* are shown in **Table S4**. The qPCR cycling conditions were as follows: 95°C for 2 min, followed by 40 cycles of 95°C for 15 s and 60°C for 1 min. The qPCR data were analyzed using the ViiA7 software, and the gene expression levels were calculated using the 2-ΔΔCT method, with GAPDH as the reference gene. The fold change in expression for each target gene between the sham and 0.5 Gy groups was calculated and plotted using PRISM 9.5.1 (GraphPad Software). Statistical analysis was performed using PRISM 9.5.1. One-way ANOVA with Bonferroni post hoc analysis was employed to determine differences between more than two groups. Values of P < 0.05 were considered statistically significant. Data are presented as means ±SEM.*P < 0.05; **P < 0.01; ***P < 0.001; P < 0.0001; n.s., non-significant.

### Bulk RNA-sequencing and analysis

Ovaries were dissected 6 hours after sham or radiation treatment with 0.5 Gy and stored in RNAlater until extraction. RNA was extracted from paired ovaries (N=6 per condition) using a miRNeasy micro extraction kit per the manufacturer’s instructions (Qiagen, 217084). RNA concentration and quality were assessed using the RNA Total RNA Nano Assay (Agilent Technologies). Libraries were constructed using the KAPA mRNA HyperPrep Kit (KAPA Biosystems). Library quality and concentration were checked using the D5000 Screen Tape (Agilent Technologies) and quantitative PCR (KAPA Biosystems). Barcoded libraries were then pooled and sequenced on the HiSeq2000 (Illumina) using TruSeq SBS Kit v4 reagents. The primary RNA-Seq processing, quality control and transcript-level quantitation, was carried out using Nextflow-based pipeline nf-core/rnaseq (version1.4.3dev) (100). Briefly, 100bp paired-end reads quality was checked using FastQC (v.0.11.9) (RRID:SCR_014583). Reads passing the quality thresholds were aligned and quantified using the Salmon quantification tool (RRID:SCR_017036) (101). Salmon v.1.3.0 was used to build an index and quantify transcript and gene expression against the reference transcriptome (Ensembl transcript release 105) using default parameters. Differential gene expression analysis was performed using the DESeq2 package (v.1.28.1) (RRID:SCR_015687) (102). Differentially expressed genes were defined as significant with an adjusted p-value (FDR) < 0.05 and Fold change ≥2.

### Single-cell RNA-seq

7-day-old wildtype and *Chek2-/-* females (N=4 per group) were exposed to sham or 0.5 Gy radiation. Pooled ovaries were dissected 6 hours after exposure and immediately dissociated into a single-cell suspension using the combined enzymatic-mechanical tissue dissociation protocol. In brief, ovaries were first incubated in collagenase IV (1mg/ml, Worthington) and DNAse (0.02%, Worthington) solution in HBSS. After 15 min incubation at 37ºC, trypsin was added to a final concentration of 0.125% and solution with ovaries was mixed by gentle pipetting. Following additional 10 min incubation at 37ºC trypsin was inactivated by adding FBS and solution with ovaries was mixed by gentle pipetting with a wide-bore tip to mechanically facilitate the dissociation of the tissue into a single-cell suspension. Cell suspensions were filtered with a 40μm mesh filter to remove debris and cell aggregates and spun down at 2000 rpm for 5 min in a swing-bucket centrifuge. Cell pellets were resuspended in 2% BSA PBS and single-cell suspensions were analyzed for viability and counted on a Countess II automated cell counter (Thermo Fisher). A total of 12,000 cells per sample were loaded onto a channel of 10X Chromium microfluidic chips for a targeted cell recovery of 6,000 cells per lane. Single-cell capture, barcoding, and library preparation were performed according to manufacturer’s protocol (10x Genomics). Sample cDNA and library quality controls were performed using the Agilent 4200 TapeStation instrument and quantified by qPCR (Kapa Biosystems/Roche). Libraries were sequenced on a NovaSeq 6000 (Illumina) with the S2 100 cycle kit targeting 50,000 reads per cell.

### Single-cell data processing and analysis

The raw sequencing reads from Illumina were aligned to the mouse reference genome mm10 using Cell Ranger V4 software (RRID:SCR_017344) with default parameters. Filtered gene counts generated after Cell Ranger were used for all the downstream analyses. The low-quality cells failing to meet the following threshold criteria were discarded, <1000 genes expressed or >20% mitochondrial transcripts or >50% ribosomal transcripts. In addition, the genes that were expressed in less than 3 cells were also discarded. The potential doublets were removed by applying the DoubletFinder (103) (RRID:SCR_018771) and DoubletDecon (104) (PMCID: PMC6983270) packages with default parameters. Only the cells marked as doublets by both the algorithms were removed. The remaining QC-pass cells were analyzed using the Seurat package (105) (RRID:SCR_016341) and batch-corrected using the Harmony package (106). In brief, the single cells were normalized based on their library size and later log-transformed. For dimensionality reduction, we applied principal component analysis on the 2000 most variable genes and used the first 30 computed PCs as an input for Leiden-based clustering. For cell type assignment, we computed the cluster-specific marker genes and visualized them in dot plots or feature plots for cell type inference. In parallel, we also used a supervised algorithm SingleR (104), that relies on pre-defined reference transcriptome profiles to compare the single-cell transcriptomes and assign the cell type labels. Finally, the single-cell results and the marker genes were visualized using either heatmaps or plots such as dot, violin, or radar plots in R language.

### Enrichment analyses of differentially expressed genes from bulk and scRNA-seq

The functional enrichment analyses were performed using g:Profiler (RRID:SCR_006809) (41) and Mus musculus as the species. Differentially expressed genes were ranked by significance (FDR) and analyzed by g:SCS multiple testing correction method and significance threshold set to 0.05. Statistical significance was calculated using a custom background list of all genes expressed in the ovary as detected by bulk RNA sequencing. Functional analysis was conducted using the Gene Ontology database for molecular function (GO:MF), cellular component (GO:CC), and biological function (GO:BF) and KEGG database for biological pathways. Minimum and Maximum term size limits were set at 10 and 1000 genes.

Gene set enrichment analysis (GSEA) for bulk RNA-seq normalized gene expression data was performed using the GSEA software (GSEA version 4.2.3) (RRID:SCR 001905) (43) and the hallmark gene set collection (RRID:SCR_016863) from the Molecular Signatures Database (MsigDB) (44). The number of permutations was set to 1000 (geneset) and dataset minimum and maximum size were set to 10 and 1000, respectively. Absolute NES (normalized enrichment score) >1.6 and FDR < 0.05 were set as cut-offs for significant enrichment. For scRNAseq data, we used the fGSEA (RRID:SCR_020938) R package for pathway enrichment analysis and tested for different gene sets like Hallmarks gene sets from MSigDB database (44) (RRID:SCR_016863). For fGSEA input, genes were pre-ranked using a fast Wilcoxon rank-sum test (presto R package V1.0.0) to test the enrichment of different gene sets. Enrichment analysis for predicted transcription factors (TF) was conducted using g:Profiler and the TRANSFAC database (with no term size limit). Further TFs enrichment analysis was performed by ChEA3 (RRID:SCR_023159) with ReMap Chip-seq library (52).

## Acknowledgments

The authors gratefully acknowledge the contribution of the Single Cell Biology Lab, Genome Technologies, Computational Sciences, and Histology Services at The Jackson Laboratory for expert assistance with the work described in this publication. These shared services are supported in part by the JAX Cancer Center (P30 CA034196). Research reported in this publication was partially supported by V Scholar Award V2017-019 and the National Cancer Institute under award number P30CA034196. The content is solely the responsibility of the authors and does not necessarily represent the official views of the NIH. The authors thank Dr. Ilya Levantis and Christina Chatzipantsiou from Lifebit for supporting Nextflow data analysis on the cloud computing platform.

## Author contributions

Conceptualization: EBF, CE, MM

Methodology: EBF, CE, MM, PK, JG

Investigation: EBF, MM, ZB, PK, CE,

Visualization: EBF, PK

Supervision: EBF

Funding acquisition: EBF

Writing—original draft: EBF

Writing—review & editing: EBF, MM, ZB, CE, PK, JG

## Competing interests

Authors declare that they have no competing interests.

## Supplementary Tables

Table S1. RRGs identified by bulk RNA-seq.

Table S2. g:Profiler analysis of RRGs.

Table S3. Gene Set Enrichment Analysis of RRGs. Table S4. Primers used in this study.

## Supplementary Data Files

Supplementary Data 1: Bulk RNAseq_Ovary_Gene_Counts_ DEGs_groupwise

Supplementary Data 2: scRNAseq_Ovary_Clusters_Marker_Genes_DEGs_groupwise

Supplementary Data 3: scRNAseq_Oocyte_Subclusters_Marker_Genes_DEGs

## Notes

### Competing Interest Statement

The authors have declared no competing interest.

