## Supplemental Figures and Tables for "Single-cell and bulk transcriptional profiling of mouse ovaries reveals novel genes and pathways associated with DNA damage response in oocytes"

**Supplementary Materials:**

**Supplementary Figures (S1-S7)**

**Supplementary Tables (S1-S4)**

### Figure S1.

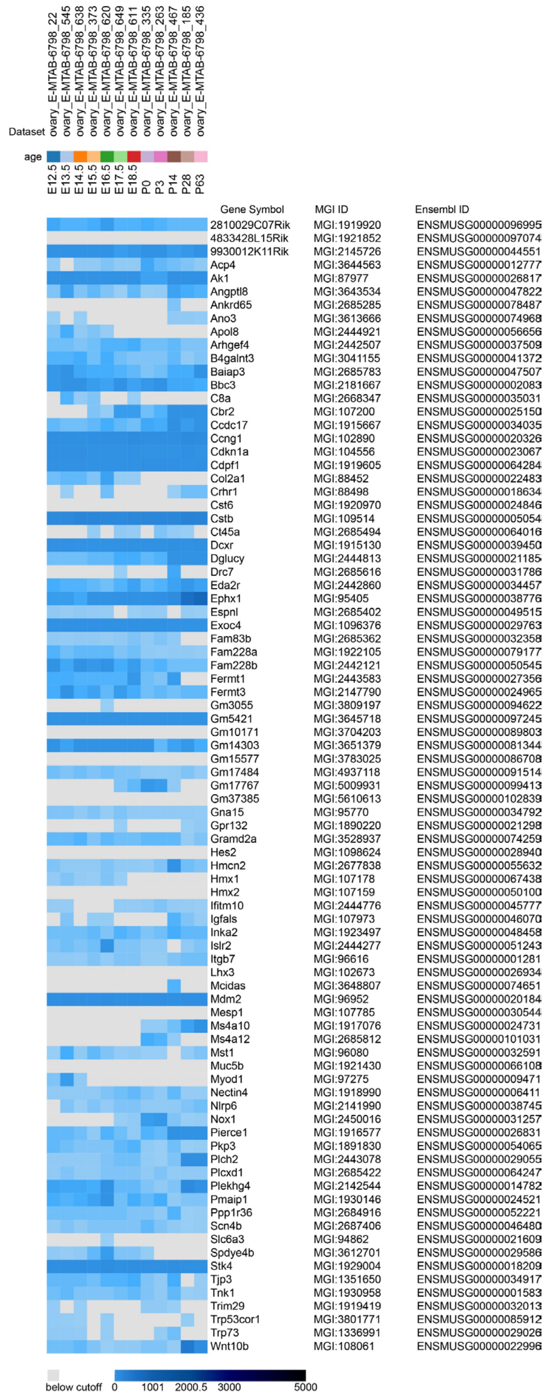


**Figure S1**. Expression of RRGs identified in the study during ovary development generated using published data and Gene Expression Database (GXD) at Mouse Genome Informatics Portal https://www.informatics.jax.org/expression.shtml

#### Figure S2.

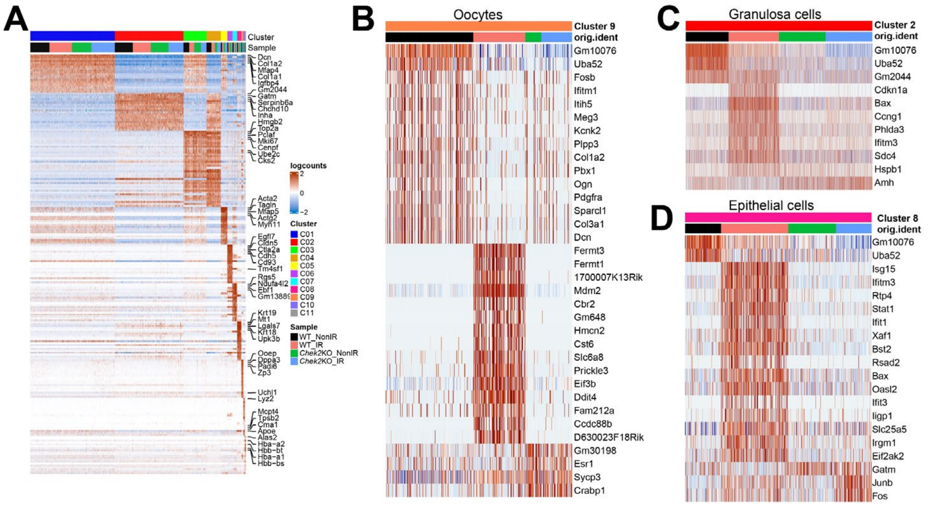


**Figure S2**. A) Heatmap of marker gene expression from single cells obtained from wildtype and *Chek2-/-* ovaries with and without radiation organized into 11 clusters. B-D) Heatmaps of differentially expressed genes in oocyte, granulosa, and epithelial cell clusters.

#### Figure S3.

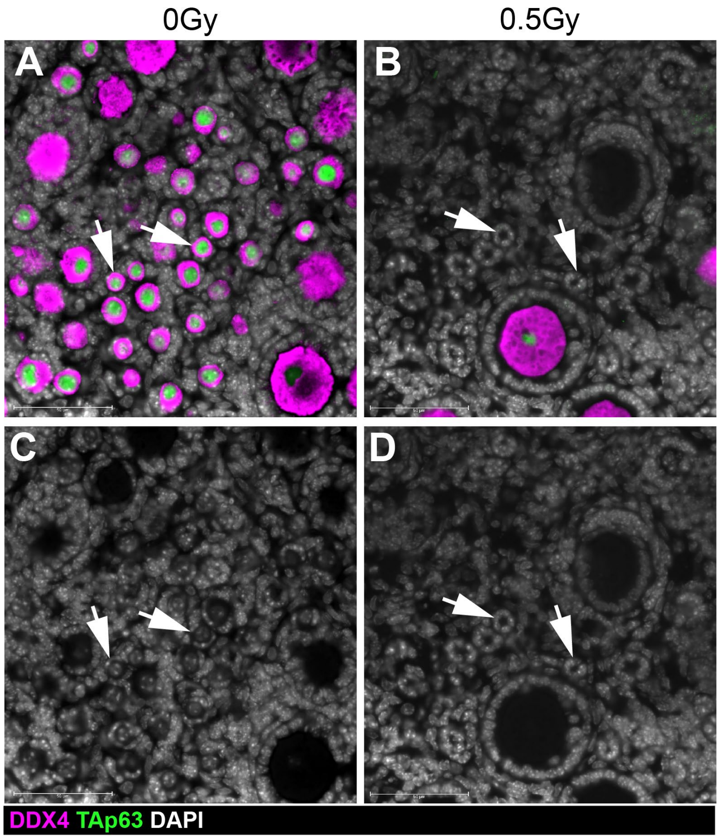


**Figure S3**. Example image of non-irradiated (A) and irradiated (B) ovaries immunostained with oocyte markers DDX4 and TAp63. C-D) Grayscale image for DAPI channel to visualize nuclei. Arrows indicate primordial follicles with an oocyte inside in non-irradiated (C) and empty follicles with visible granulosa cells but without the oocyte in the irradiated ovary (D).

### Figure S4.

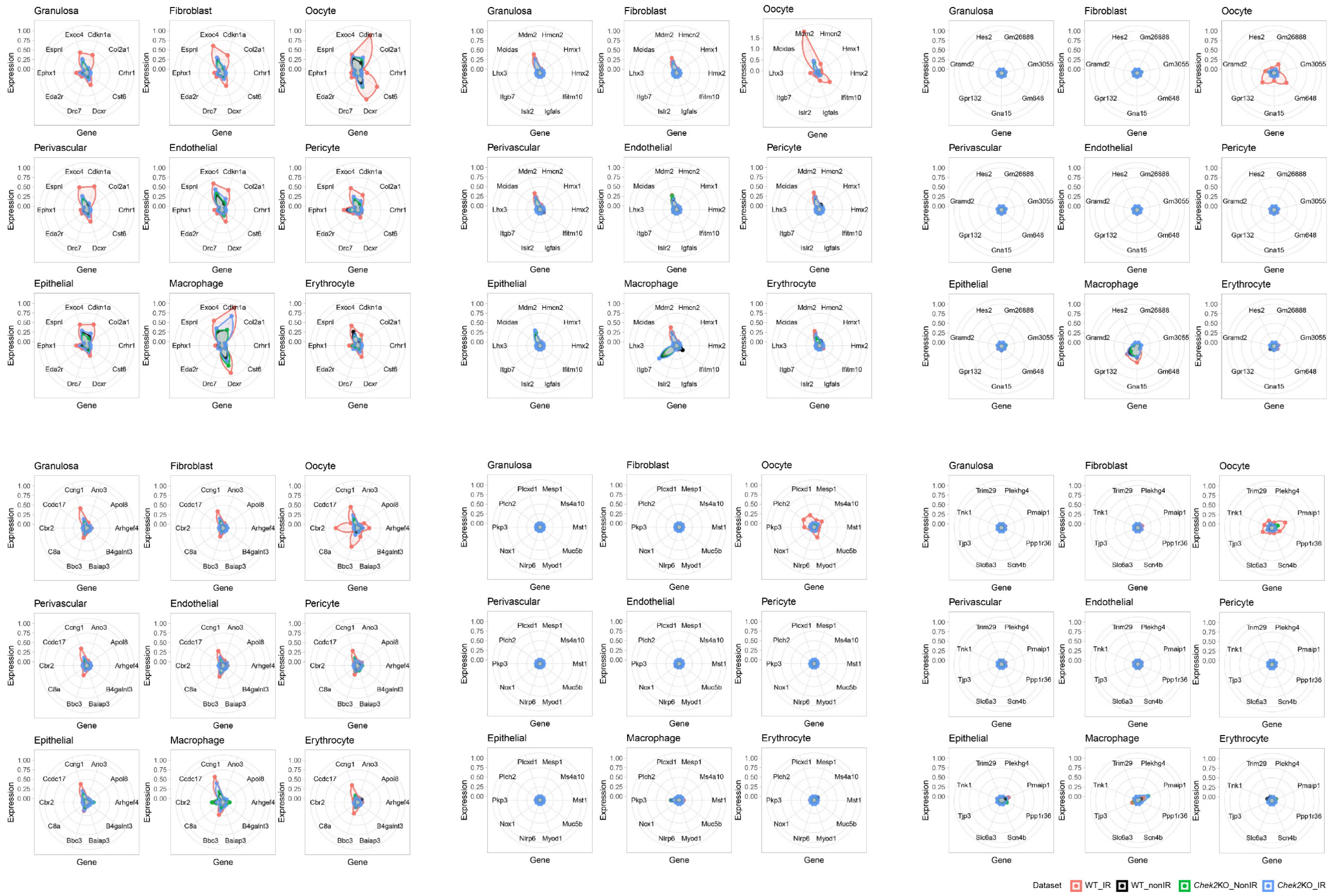


**Figure S4**. Radar charts showing expression of RRGs in different cell types.

#### Figure S5.

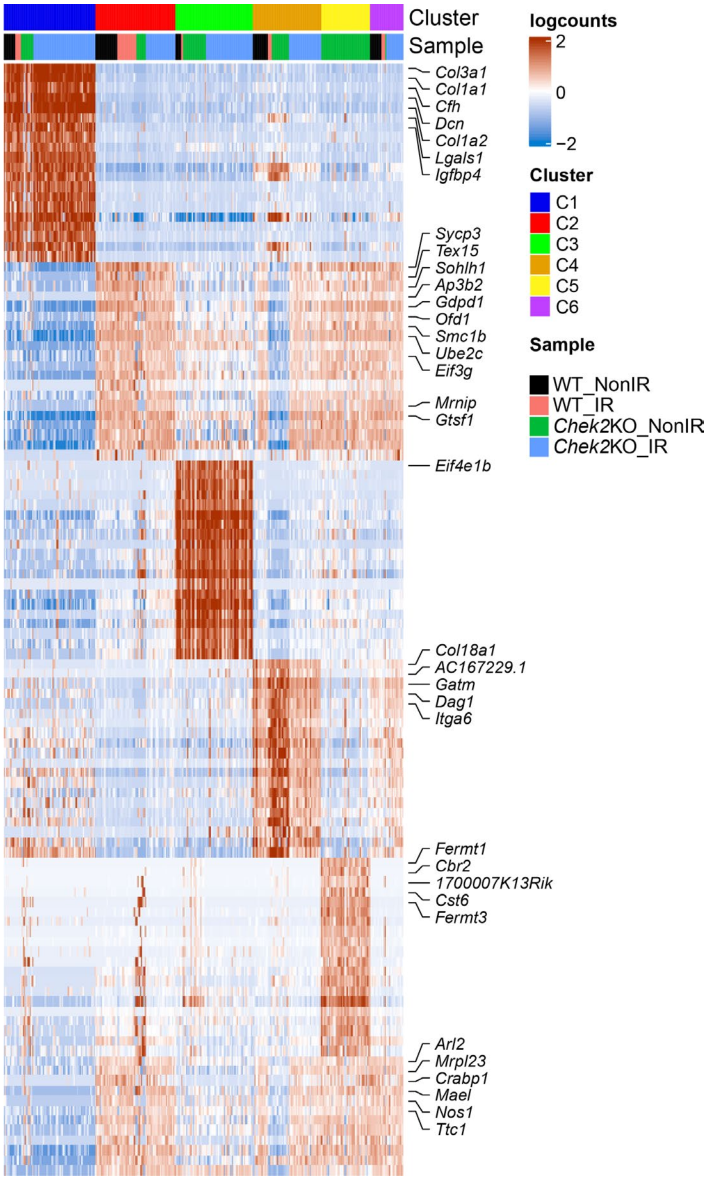


**Figure S5.** A) Heatmap of marker gene expression for six oocyte subclusters identified by de novo re-clustering.

#### Figure S6.

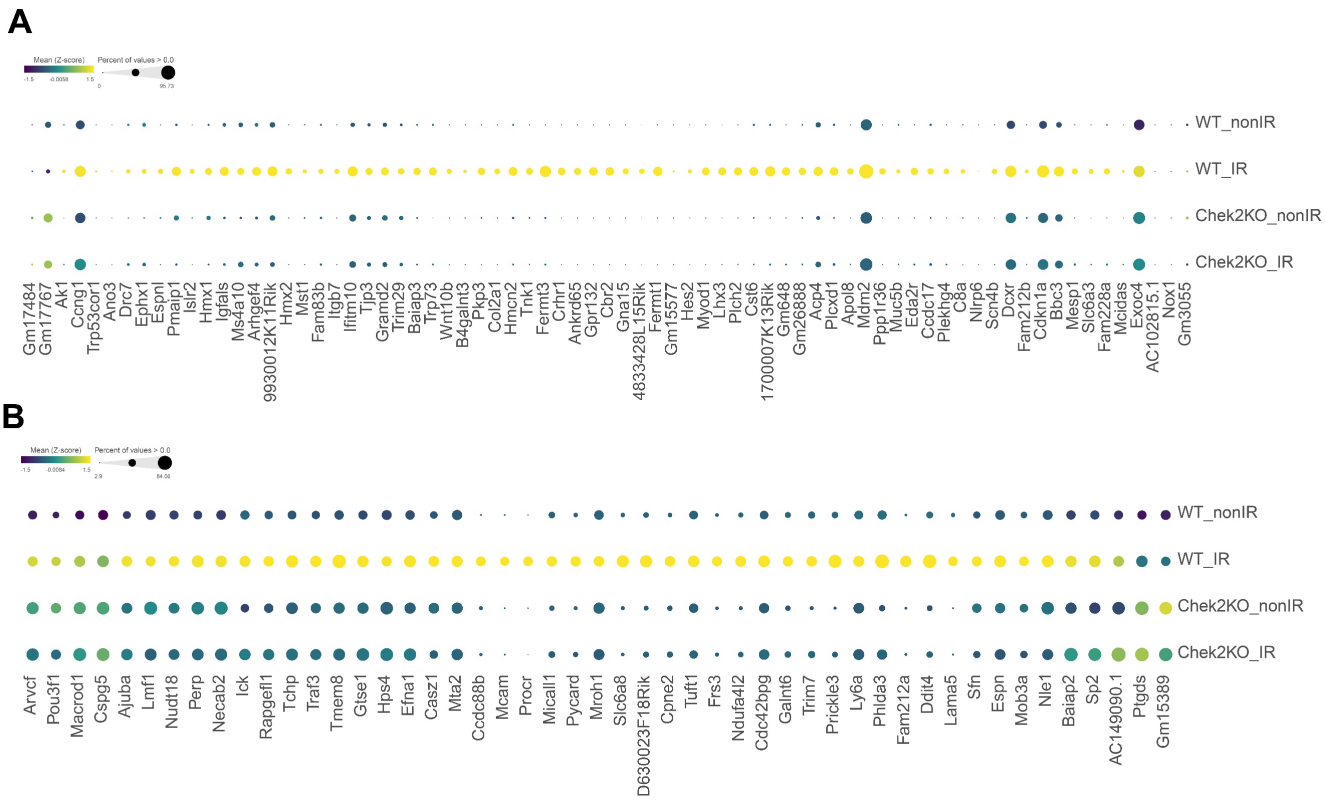


**Figure S6.** Dot plot showing expression of RRGs from bulk (A) and additional RRGs from subcluster analysis (B) in all oocytes from wildtype and *Chek2-/-* ovaries with and without radiation.

#### Figure S7.

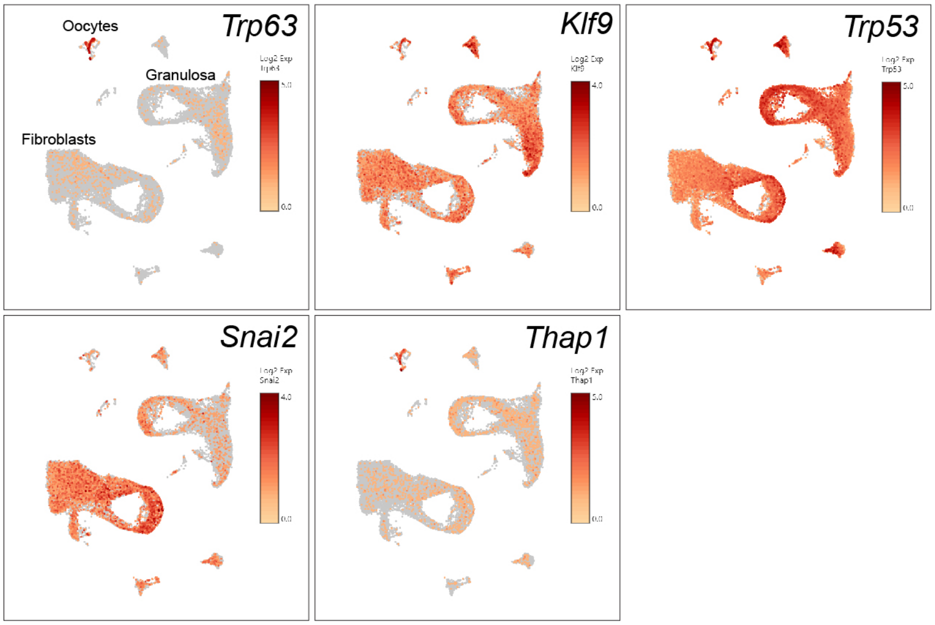


**Figure S7.** Gene expression levels of transcription factors predicted to regulate RRGs overlaid on UMAP plots with the oocyte, granulosa, and fibroblast clusters indicated.

#### Supplementary Tables:

### Supplementary Table S1. RRGs identified by bulk RNA-seq.

**Table S1. Radiation Responsive Genes in the ovary**
Part 1. Response in wildtype ovaries.

| MGI Gene/ Marker ID | Symbol | LogFC WT | Adj p-value | Feature Type | Gene Ontology terms |
|---|---|---|---|---|---|
| **MGI:5610613** | *Gm37385* | 22.7 | 8.77E-05 | unclassified gene | |
| **MGI:3809197** | *Gm3055* | 20.6 | 1.02E-03 | protein coding gene | |
| **MGI:102673** | *Lhx3* | 7.8 | 1.14E-10 | protein coding gene | cell differentiation; DNA-binding transcription factor activity, RNA polymerase II-specific |
| **MGI:1921852** | *4833428L15Rik* | 6.9 | 1.06E-07 | lncRNA gene | |
| **MGI:2685285** | *Ankrd65* | 6.7 | 1.22E-18 | protein coding gene | |
| **MGI:107159** | *Hmx2* | 6.5 | 3.27E-05 | protein coding gene | cell differentiation; DNA-binding transcription factor activity, RNA polymerase II-specific |
| **MGI:2444921** | *Apol8* | 5.8 | 6.63E-08 | protein coding gene | chloride channel activity; extracellular space; high-density lipoprotein particle; intracellular membrane-bounded organelle |
| **No associated gene** | *Gm43301* | 5.7 | 1.22E-04 | lncRNA gene | |
| **MGI:3613666** | *Ano3* | 5.6 | 1.27E-30 | protein coding gene | calcium activated phospholipid scrambling; chloride channel activity; chloride transmembrane transport |
| **MGI:3783025** | *Gm15577* | 5.5 | 5.92E-03 | lncRNA gene | |
| **MGI:107785** | *Mesp1* | 5.5 | 2.99E-03 | protein coding gene | DNA-binding transcription activator activity, RNA polymerase II-specific; Notch signaling pathway |
| **MGI:1098624** | *Hes2* | 5.5 | 6.45E-08 | protein coding gene | DNA-binding transcription factor activity, RNA polymerase II-specific; DNA-binding transcription repressor activity, RNA polymerase II-specific |
| **MGI:2443078** | *Plch2* | 5.3 | 8.57E-51 | protein coding gene | calcium ion binding; intracellular signal transduction; phosphatidylinositol-mediated signaling; phospholipase C activity; plasma membrane; signal transduction |
| **MGI:94862** | *Slc6a3* | 5.2 | 2.50E-11 | protein coding gene | cell surface; dopamine transport; integral component of membrane; signaling receptor binding |
| **MGI:107200** | *Cbr2* | 5.1 | 2.91E-05 | protein coding gene | carbonyl reductase (NADPH) activity; glucose metabolic process; mitochondrion; oxidoreductase activity |
| **MGI:97275** | *Myod1* | 4.8 | 6.15E-07 | protein coding gene | cell differentiation; cellular response to estradiol stimulus; cellular response to oxygen levels; cellular response to tumor necrosis factor; DNA-binding transcription activator activity, RNA polymerase II-specific; ubiquitin protein ligase binding |
| **MGI:88498** | *Crhr1* | 4.7 | 2.07E-09 | protein coding gene | adenylate cyclase-activating G protein-coupled receptor; signaling pathway; cell surface receptor signaling pathway; endosome; transmembrane signaling receptor activity; vesicle |
| **MGI:2685402** | *Espnl* | 4.7 | 2.42E-18 | protein coding gene | actin binding; cell projection; stereocilium tip; stereocilium tip |
| **MGI:2668347** | *C8a* | 4.7 | 1.39E-03 | protein coding gene | complement activation; cytolysis; immune response; membrane attack complex |
| **MGI:2443583** | *Fermt1* | 4.5 | 9.82E-06 | protein coding gene | cell adhesion; cell junction; focal adhesion; integrin-mediated signaling pathway; negative regulation of; canonical Wnt signaling pathway; positive regulation of cell adhesion mediated by integrin |
| **MGI:1890220** | *Gpr132* | 4.5 | 1.42E-15 | protein coding gene | G1/S transition of mitotic cell cycle; G protein-coupled receptor signaling pathway; negative regulation of G2/M; transition of mitotic cell cycle; plasma membrane; signal transduction |
| **MGI:2685616** | *Drc7* | 4.4 | 5.93E-07 | protein coding gene | cell differentiation; cell projection; cilium |
| **MGI:88452** | *Col2a1* | 4.3 | 1.35E-06 | protein coding gene | cellular response to BMP stimulus; extracellular region |
| **MGI:3648807** | *Mcidas* | 4.2 | 2.60E-02 | protein coding gene | cell cycle; negative regulation of cell cycle; negative regulation of DNA replication |
| **MGI:2444776** | *Ifitm10* | 4.2 | 5.11E-37 | protein coding gene | integral component of membrane; membrane |
| **MGI:1916577** | *Pierce1 (1700007K13Rik )* | 4.1 | 1.10E-18 | protein coding gene | cellular response to DNA damage stimulus; regulation of gene expression |
| **MGI:2685494** | *Gm648* | 4.1 | 2.26E-10 | protein coding gene | |
| **MGI:95770** | *Gna15* | 4.0 | 3.05E-05 | protein coding gene | adenylate cyclase-modulating G protein-coupled receptor signaling pathway; GTPase activity membrane; phospholipase C-activating G protein-coupled receptor signaling pathway |
| **MGI:1918990** | *Nectin4 (Pvrl4)* | 3.9 | 3.12E-10 | protein coding gene | cell junction |
| **MGI:1919920** | *2810029C07Rik* | 3.9 | 9.10E-37 | lncRNA gene | |
| **MGI:2444277** | *Islr2* | 3.8 | 2.02E-11 | protein coding gene | cell surface; integral component of membrane; protein binding |
| **MGI:2687406** | *Scn4b* | 3.7 | 2.07E-11 | protein coding gene | ion transport; plasma membrane; sodium channel activity |
| **MGI:2684916** | *Ppp1r36* | 3.7 | 1.11E-09 | protein coding gene | negative regulation of phosphatase activity; phosphatase binding |
| **MGI:3612701** | *Spdye4b (4933411G11Rik )* | 3.7 | 3.15E-03 | protein coding gene | protein kinase binding |
| **MGI:2147790** | *Fermt3* | 3.6 | 4.15E-10 | protein coding gene | cell adhesion; cell junction; integrin-mediated signaling pathway |
| **MGI:3645718** | *Gm5421* | 3.6 | 1.67E-02 | pseudogene | |
| **MGI:2685362** | *Fam83b* | 3.3 | 3.85E-23 | protein coding gene | phosphatidylinositol 3-kinase catalytic subunit binding; signal transduction |
| **MGI:1921430** | *Muc5b* | 3.3 | 4.79E-08 | protein coding gene | extracellular matrix; regulation of macrophage activation |
| **MGI:2444813** | *Dglucy (9030617O03Rik )* | 3.3 | 5.40E-16 | protein coding gene | D-glutamate cyclase activity; lyase activity; mitochondrion |
| **MGI:107178** | *Hmx1* | 3.2 | 2.56E-05 | protein coding gene | DNA-binding transcription factor activity, RNA polymerase II-specific |
| **MGI:3801771** | *Trp53cor1* | 3.2 | 2.03E-04 | lncRNA gene | negative regulation of gene expression; ribonucleoprotein complex |
| **MGI:1920970** | *Cst6* | 3.2 | 4.08E-04 | protein coding gene | epidermis development |
| **MGI:1930958** | *Tnk1* | 3.0 | 7.58E-06 | protein coding gene | ATP binding; kinase activity; negative regulation of Ras protein signal transduction; transmembrane receptor protein tyrosine kinase signaling pathway |
| **MGI:108061** | *Wnt10b* | 3.0 | 1.04E-02 | protein coding gene | canonical Wnt signaling pathway; G2/M transition of mitotic cell cycle; negative regulation of transcription by RNA polymerase II; regulation of cell cycle |
| **MGI:96080** | *Mst1* | 3.0 | 4.59E-04 | protein coding gene | extracellular region; negative regulation of epithelial cell apoptotic process; receptor tyrosine kinase binding; regulation of receptor signaling pathway via JAK-STAT serine-type endopeptidase activity |
| **MGI:1915667** | *Ccdc17* | 3.0 | 7.06E-18 | protein coding gene | |
| **MGI:5477382** | *Gm26888* | 2.8 | 4.60E-09 | lncRNA gene | |
| **MGI:1336991** | *Trp73* | 2.8 | 8.25E-11 | protein coding gene | apoptotic process; cell cycle; cell junction; cellular response to DNA damage stimulus; DNA-binding transcription activator activity, RNA polymerase II-specific; intrinsic apoptotic signaling pathway in response to DNA damage by p53 class mediator; MDM2/MDM4 family protein binding; negative regulation of JUN kinase activity; p53 binding; positive regulation of apoptotic process; positive regulation of MAPK cascade |
| **MGI:1930146** | *Pmaip1* | 2.7 | 1.16E-16 | protein coding gene | activation of cysteine-type endopeptidase activity; involved in apoptotic process; positive regulation of apoptotic process; positive regulation of DNA damage response, signal transduction by p53 class mediator; reactive oxygen species metabolic process; regulation of apoptotic process |
| **MGI:3643534** | *Angptl8 (Gm6484)* | 2.6 | 3.32E-04 | protein coding gene | extracellular region; negative regulation of lipoprotein lipase activity |
| **MGI:1891830** | *Pkp3* | 2.6 | 6.08E-07 | protein coding gene | adherens junction; alpha-catenin binding; cell-cell adhesion; cell-cell junction assembly |
| **MGI:2142544** | *Plekhg4* | 2.6 | 6.96E-03 | protein coding gene | activation of GTPase activity |
| **MGI:1351650** | *Tjp3* | 2.5 | 4.90E-06 | protein coding gene | cell-cell adhesion; cell-cell junction; cell surface; regulation of G1/S transition of mitotic cell cycle |
| **MGI:3644563** | *Acp4 (Acpt)* | 2.5 | 1.09E-07 | protein coding gene | acid phosphatase activity; negative regulation of ERBB4 signaling pathway; peptidyl-tyrosine dephosphorylation involved in inactivation of protein kinase activity; protein tyrosine phosphatase activity |

| | | | | | |
|---|---|---|---|---|---|
| **MGI:96616** | *Itgb7* | 2.4 | 2.61E-04 | protein coding gene | cell adhesion; cell adhesion mediated by integrin; focal adhesion; integrin-mediated signaling pathway |
| **MGI:1919419** | *Trim29* | 2.4 | 5.51E-03 | protein coding gene | innate immune response; lysosome; negative regulation of transcription by RNA polymerase II; p53 binding |
| **MGI:1922105** | *Fam228a* | 2.4 | 1.30E-02 | protein coding gene | |
| **MGI:3528937** | *Gramd2* | 2.3 | 5.77E-08 | protein coding gene | extrinsic component of cytoplasmic side of plasma membrane; integral component of membrane; phosphatidylinositol binding |
| **MGI:1917076** | *Ms4a10* | 2.1 | 2.42E-03 | protein coding gene | cell surface receptor signaling pathway; integral component of membrane; membrane |
| **MGI:4937118** | *Gm17484* | 2.1 | 5.54E-05 | lncRNA gene | |
| **MGI:2145726** | *9930012K11Rik* | 2.1 | 1.26E-12 | protein coding gene | |
| **MGI:102890** | *Ccng1* | 2.0 | 5.64E-105 | protein coding gene | cell cycle; cell division; cyclin-dependent protein; serine/threonine kinase regulator activity; cytoplasm; mitotic G2 DNA damage checkpoint signaling; negative regulation of apoptotic process |
| **MGI:87977** | *Ak1* | 2.0 | 1.64E-05 | protein coding gene | adenylate kinase activity; regulation of G1/S transition of mitotic cell cycle |
| **MGI:2685783** | *Baiap3* | 2.0 | 1.95E-05 | protein coding gene | calcium ion binding; G protein-coupled receptor signaling pathway; late endosome membrane; regulation of dense core granule exocytosis |
| **MGI:2685422** | *Plcxd1* | 1.9 | 1.10E-07 | protein coding gene | lipid metabolic process |
| **MGI:2181667** | *Bbc3* | 1.9 | 3.41E-11 | protein coding gene | activation of cysteine-type endopeptidase activity; involved in apoptotic process; cellular response to DNA damage stimulus; cellular response to ionizing radiation; intrinsic apoptotic signaling pathway; intrinsic apoptotic signaling pathway by p53 class mediator; PUMA-BCL-xl complex; response to endoplasmic reticulum stress |
| **MGI:107973** | *Igfals* | 1.9 | 2.92E-02 | protein coding gene | cell adhesion; extracellular region |
| **MGI:2677838** | *Hmcn2* | 1.8 | 1.92E-03 | protein coding gene | basement membrane; cell junction; collagen-containing extracellular matrix; response to stimulus |
| **MGI:1923497** | *Inka2 (Fam212b)* | 1.8 | 2.95E-13 | protein coding gene | negative regulation of catalytic activity; protein serine/threonine kinase inhibitor activity |
| **MGI:2442860** | *Eda2r* | 1.8 | 2.19E-21 | protein coding gene | cell differentiation; integral component of membrane; intrinsic apoptotic signaling pathway by p53 class mediator; positive regulation of I-kappaB kinase/NF-kappaB signaling; positive regulation of JNK cascade<br>programmed cell death; tumor necrosis factor-mediated signaling pathway |
| **MGI:3041155** | *B4galnt3* | 1.7 | 3.26E-03 | protein coding gene | acetylgalactosaminyltransferase activity; glycosyltransferase activity; Golgi apparatus |
| **MGI:95405** | *Ephx1* | 1.7 | 3.26E-03 | protein coding gene | integral component of membrane; plasma membrane; response to toxic substance |
| **MGI:104556** | *Cdkn1a* | 1.7 | 5.54E-05 | protein coding gene | cell cycle; cellular response to DNA damage stimulus; cellular response to ionizing radiation; cellular response to UV-B; cyclin-dependent protein serine/threonine kinase inhibitor activity; DNA damage response, signal; transduction by p53 class mediator resulting in cell cycle arrest; mitotic G2 DNA damage checkpoint signaling; PCNA-p21 complex; regulation of cell cycle G1/S phase transition |
| **MGI:1915130** | *Dcxr* | 1.6 | 2.91E-03 | protein coding gene | carbonyl reductase (NADPH) activity; L-xylulose reductase (NADP+) activity; membrane; oxidoreductase activity; positive regulation of reactive oxygen species; metabolic process |
| **MGI:2442507** | *Arhgef4* | 1.5 | 6.22E-04 | protein coding gene | intracellular signal transduction; lamellipodium assembly; protein domain specific binding |
| **MGI:1096376** | *Exoc4* | 1.4 | 9.61E-06 | protein coding gene | cell projection; exocytosis; regulation of protein transport; small GTPase binding; vesicle tethering involved in exocytosis |
| **MGI:96952** | *Mdm2* | 1.3 | 3.40E-07 | protein coding gene | apoptotic process; cellular response to gamma radiation; DNA damage response, signal transduction by p53 class mediator resulting in cell cycle arrest; NEDD8 ligase activity; p53 binding |
| **MGI:5009931** | *Gm17767* | -2.2 | 4.93E-03 | lncRNA gene | |
| **MGI:2450016** | *Nox1* | -2.6 | 7.06E-05 | protein coding gene | cell junction; cellular response to hyperoxia; endosome; extracellular matrix organization; NADPH oxidase complex; oxidoreductase activity; positive regulation of JNK cascade; positive regulation of MAPK cascade; positive regulation of oxidative stress-induced intrinsic; apoptotic signaling pathway; small GTPase binding |
| **MGI:2141990** | *Nlrp6* | -3.1 | 8.69E-03 | protein coding gene | activation of cysteine-type endopeptidase activity; acute inflammatory response; ATP binding; inflammasome complex; necroptotic process; negative regulation of ERK1 and ERK2 cascade; negative regulation of I-kappaB kinase/NF-kappaB signaling; NLRP6 inflammasome complex |
| **No associated gene** | *AC102815.1* | -15.2 | 1.96E-02 | unclassified gene | |
| **MGI:3704203** | *Gm10171* | -20.2 | 1.64E-07 | pseudogene | |
| **MGI:3651379** | *Gm14303* | -22.2 | 1.71E-04 | pseudogene | |

| Part 2. Response in *Chek2-/-* ovaries. | | | | | |
|---|---|---|---|---|---|
| **MGI Gene/ Marker ID** | **Symbol** | **LogFC *Chek2-/-*** | **Adj p-value** | **Feature Type** | **Gene Ontology terms** |
| **MGI:5610613** | *Gm37385* | 26.4 | 1.77E-05 | unclassified gene | |
| **MGI:1338893** | *Padi1* | -6.7 | 0.032901 | protein coding gene | calcium ion binding; hydrolase activity; protein-arginine deiminase activity |
| **MGI:102673** | *Gm43520* | -22.9 | 1.77E-05 | unclassified gene | |

#### Supplementary Table S2. Gene enrichment analysis of RRGs with g:Profiler.

| Source | Term Name | Term ID | Adj.pvalue | -log10 adj.pvalue | Genes |
|---|---|---|---|---|---|
| **GO:BP** | intrinsic apoptotic signaling pathway by p53 class mediator | GO:0072332 | 0.001 | 3.15 | *Eda2r, Pmaip1, Bbc3, Trp73, Mdm2, Cdkn1a* |
| **GO:BP** | signal transduction by p53 class mediator | GO:0072331 | 0.019 | 1.71 | *Eda2r, Pmaip1, Bbc3, Trp73, Mdm2, Cdkn1a* |
| **KEGG** | p53 signaling pathway | KEGG:04115 | 0.000 | 6.62 | *Ccng1, Pmaip1, Bbc3, Trp73, Mdm2, Cdkn1a* |
| **KEGG** | Platinum drug resistance | KEGG:01524 | 0.001 | 3.05 | *Pmaip1, Bbc3, Mdm2, Cdkn1a* |
| **KEGG** | Apoptosis - multiple species | KEGG:04215 | 0.013 | 1.88 | *Pmaip1, Bbc3* |
| **KEGG** | Human papillomavirus infection | KEGG:05165 | 0.024 | 1.62 | *Hes2, Mdm2, Col2a1, Cdkn1a, Itgb7, Wnt10b* |
| **KEGG** | Colorectal cancer | KEGG:05210 | 0.036 | 1.44 | *Pmaip1, Bbc3, Cdkn1a* |
| **WP** | p53 signaling | WP:WP2902 | 0.000 | 7.07 | *Ccng1, Pmaip1, Bbc3, Trp73, Mdm2, Cdkn1a* |
| **WP** | Apoptosis | WP:WP1254 | 0.004 | 2.41 | *Pmaip1, Trp73, Mdm2* |
| **WP** | Hypoxia-dependent self-renewal of myoblasts | WP:WP5023 | 0.009 | 2.04 | *Myod1, Cdkn1a* |
| **MIRNA** | mmu-miR-23a-3p | MIRNA:mmu-miR-23a-3p | 0.030 | 1.52 | *Pmaip1, Bbc3* |

**Supplementary Table S3. Gene Set Enrichment Analysis of RRGs (GSEA).**

| HALLMARK PATHWAY | ES | NES | NOM p-val | FDR q-val | TOP GENES (HUMAN SYMBOLS) |
|---|---|---|---|---|---|
| **P53 PATHWAY** | 0.67 | 2.20 | 0.00 | 0.00 | *CCNG1, AK1, CLCA2, CDKN1A, DCXR, EPHX1, MDM2, PIDD1, PHLDA3, BTG2, ZMAT3, ZNF365, GLS2, MAPKAPK3, EPHA2, SFN, BAX, SLC19A2, DEF6, DDIT4* |
| **INTERFERON ALPHA RESPONSE** | 0.67 | 2.05 | 0.00 | 0.00 | *DHX58, RTP4, TRIM14, IRF9, LGALS3BP, TMEM140, TAP1, EIF2AK2, SAMD9L, PARP9, IRF7, HELZ2, PROCR, DDX60, TRIM21, ISG15, PSMB8, IFITM3, GBP4, HERC6* |
| **INTERFERON GAMMA RESPONSE** | 0.57 | 1.86 | 0.00 | 0.00 | *CDKN1A, ITGB7, DHX58, OAS2, RTP4, TRIM14, IRF9, LGALS3BP, TAP1, EIF2AK2, SAMD9L, DDX58, XAF1, STAT1, IRF7, HELZ2, DDX60, CSF2RB, TRIM21, ISG15* |
| **APOPTOSIS** | 0.53 | 1.70 | 0.00 | 0.01 | *PMAIP1, GNA15, CDKN1A, BTG2, BAX, TAP1, CD14, GUCY2D, IFITM3, CASP7, MMP2, GADD45A, IER3, CASP1, PLCB2, GSR, FAS, HGF, DPYD, TNFSF10* |
| **TNFA SIGNALING VIA NFKB** | 0.51 | 1.69 | 0.00 | 0.01 | *CDKN1A, BTG2, TAP1, EGR2, KYNU, DDX58, EGR1, DRAM1, LIF, F3, DUSP2, PLEK, PTGER4, EFNA1, IFIH1, LAMB3, SOCS3, IL7R, FOS, PLK2* |
| **INFLAMMATORY RESPONSE** | 0.49 | 1.61 | 0.00 | 0.02 | *GPR132, GNA15, CDKN1A, BTG2, RTP4, EIF2AK2, IRF7, LIF, RNF144B, BEST1, F3, CD14, ADORA2B, PTGER4, P2RX7, SEMA4D, HRH1, IL7R, ITGB8, CD48* |
| **IL6 JAK STAT3 SIGNALING** | 0.52 | 1.56 | 0.01 | 0.03 | *IRF9, STAT1, IL1R2, CSF2RB, CD14, CCR1, SOCS3, CXCL9, DNTT, IL18R1, SOCS1, STAT2, CSF3R, FAS, PIK3R5, EBI3, MAP3K8, ACVR1B, PF4, CD44* |
| **APICAL JUNCTION** | 0.47 | 1.54 | 0.00 | 0.03 | *EXOC4, NECTIN4, COL17A1, NFASC, MAP3K20, CLDN4, ACTA1, PPP2R2C, CDH4, PARD6G, CLDN14, LIMA1, GRB7, VCL, BAIAP2, ICAM5, THY1, MAP4K2, CDH3, LAMB3* |
| **ESTROGEN RESPONSE LATE** | 0.47 | 1.53 | 0.00 | 0.03 | *PKP3, DCXR, TRIM29, TJP3, RAPGEFL1, OVOL2, SFN, ASS1, CLIC3, ST14, CA2, CACNA2D2, TOB1, SCNN1A, MAPT, PERP, DUSP2, TFAP2C, CELSR2, PGR* |
| **COMPLEMENT** | 0.46 | 1.52 | 0.00 | 0.03 | *TMPRSS6, ADRA2B, CDK5R1, CA2, KLK1, MMP15, KYNU, IRF7, F3, PLEK, CASP7, PSMB9, GCA, LCP2, F7, ITIH1, CFB, ACTN2, ITGAM, TFPI2* |
| **COAGULATION** | 0.46 | 1.47 | 0.02 | 0.04 | *C8A, MST1, TMPRSS6, MMP15, F3, PLEK, CTSE, MMP2, ITIH1, F13B, HPN, CFB, P2RY1, TFPI2, C1QA, F10, KLK8, LEFTY2, WDR1, GP9* |
| **ESTROGEN RESPONSE EARLY** | 0.44 | 1.47 | 0.01 | 0.05 | *PMAIP1, ESRP2, TJP3, RAPGEFL1, OVOL2, SFN, CLIC3, SLC19A2, SYT12, TOB1, SCNN1A, MAPT, ELF3, TFAP2C, CELSR2, ADCY1, PGR, TTC39A, MUC1, FOS* |

ES- Enrichment Score; NES- Normalized Enrichment Score; NOM p-val- Nominal pvalue; FDR q-val- False discovery rate

**Supplementary Table S4. Primers used in this study.**

| **RT-qPCR Primers** | | |
|---|---|---|
| **Gene** | **Sequence** | **Direction** |
| *Ankrd65* | AGTGGCTAAGGGCATTGAAATA | Forward |
| *Ankrd65* | GGAGATGGCTGACAACTTCTAC | Reverse |
| *Cbr2* | GGTAGCCAGGGACATGATTAAC | Forward |
| *Cbr2* | GAGCTGTAGGTGATCAAGTTAGG | Reverse |
| *Cdkn1a* | TTAGGCAGCTCCAGTGGCAACC | Forward |
| *Cdkn1a* | ACCCCCACCACCACACACCATA | Reverse |
| *Fermt1* | CATGCAAATGGAGAGCAGCAG | Forward |
| *Fermt1* | TTCCCACCACAGAGCATAGTC | Reverse |
| *Fermt3* | ATGGAGGCTCAGGGAACAAA | Forward |
| *Fermt3* | CTTGGCCTTGAACTTTCGCT | Reverse |
| *Gapdh* | TCCATGACAACTTTGGCATTG | Forward |
| *Gapdh* | CAGTCTTCTGGGTGGCAGTGA | Reverse |
| *Ddx4* | GTGGAAATACTGGCAGAGCG | Forward |
| *Ddx4* | ATCCTGTTGAGCGTCTGACA | Reverse |

| **Genotyping primers** | | | |
|---|---|---|---|
| **Gene** | **Probe** | **Sequence** | **Direction** |
| *Chek2* | *Wildtype* | CTTGTCCTGCTGGACTCACA | Forward |
| *Chek2* | *Wildtype* | GCAACCGTTACCTACCCTGA | Reverse |

| | | | |
|---|---|---|---|
| *Chek2* | *Mutant* | CGGTCGCTACCATTACCAGT | Forward |
| *Chek2* | *Mutant* | CAGCGCTTATCCCAACACT | Reverse |
| *Trp53* | *Wildtype* | CAGCCTCTGTTCCACATACACT | Forward |
| *Trp53* | *Mutant* | AGGCTTAGAGGTGCAAGCTG | Forward |
| *Trp53* | *Common* | TGGATGGTGGTATACTCAGAGC | Reverse |
| *Trp63* | *Common* | ACCTGGCTTCCTTCTCATTG | Forward |
| *Trp63* | *Common* | CTTTGATACGCTGCTGCTTG | Reverse |

#### Supplementary Data Files.

Supplementary Data 1: Bulk RNAseq_Ovary_Gene_Counts_ DEGs_groupwise
Supplementary Data 2: scRNAseq_Ovary_Clusters_Marker_Genes_DEGs_groupwise
Supplementary Data 3: scRNAseq_Oocyte_Subclusters_Marker_Genes_DEGs
